# Environment-mediated interactions cause an externalized and collective memory in microbes

**DOI:** 10.1101/2024.09.09.612037

**Authors:** Shubham Gajrani, Xiaozhou Ye, Christoph Ratzke

## Abstract

Microbes usually live in complex communities interacting with many other microbial species. These interactions determine who can persist in a community and how the overall community forms and functions. Microbes often exert interactions by chemically changing the environment, like taking up nutrients or producing toxins. These environmental changes can persist over time. We show here that such lasting environmental changes can cause a memory effect where current growth conditions alter interaction outcomes in the future. Importantly, this memory is only stored in the environment and not inside the bacterial cells. Only the collective effort of many bacteria can build up this memory, making it an emergent property of bacterial populations. This “population memory” can also impact the assembly of more complex communities and lead to different final communities depending on the system’s past. Overall, we show that to understand interaction outcomes fully, we not only have to consider the interacting species and abiotic conditions but also the system’s history.

## Main

Information storage is a central ability of living systems. Usually, memory is stored in an internal physical storage unit like the brain in higher organisms^1^. But, also on the much smaller scale of a single cell, information can be stored, e.g., by modification of the genome^2,3^, positive feedback loops of gene expression^4^, or inheritance of long-lived proteins^5^. In particular, microbes can possess cellular memory for the timing of cellular decision-making^6^, sporulation^7^, or prolonged persistence to antibiotics^8^.

In contrast, several examples of information stored outside the organism as externalized memory exist. Thus, social insects can deposit pheromone trails that guide them and their fellows^9^. Recently, the slime mold *Physarum polycephalum* rose to fame, solving a complex spatial task by depositing slime as a clue of its former location—it uses this slime as an externalized memory ^10,11^.

Also, microbes are known for secreting many metabolites, from simple byproducts of energy metabolism to complex toxins^12^. The metabolites often directly impact their and other microbes’ growth and thus can drive microbial interactions^13,14^. Which metabolites the bacteria secrete depends strongly on the microbial metabolism and, therefore, the environmental conditions the microbes face, such as temperature^15^, oxygen levels^16^, or toxins concentration^17^. This situation raises the question of whether microbes can also show a type of externalized memory where the current conditions lead to different chemical changes in the environment that further impact the growth and interactions of microbes in the future (Fig. 1A). Since a single microbe has a minimal potential to change the environment^18,19^, we hypothesize that the formation of externalized memory should require the collective action of a large number of microbes. Microbial communities could establish a memory by collectively secreting metabolites despite no memory being present in the individual cell (Fig. 1A). Such a memory would be an emergent collective phenomenon, and we call it population memory accordingly.

**Fig. 1:**
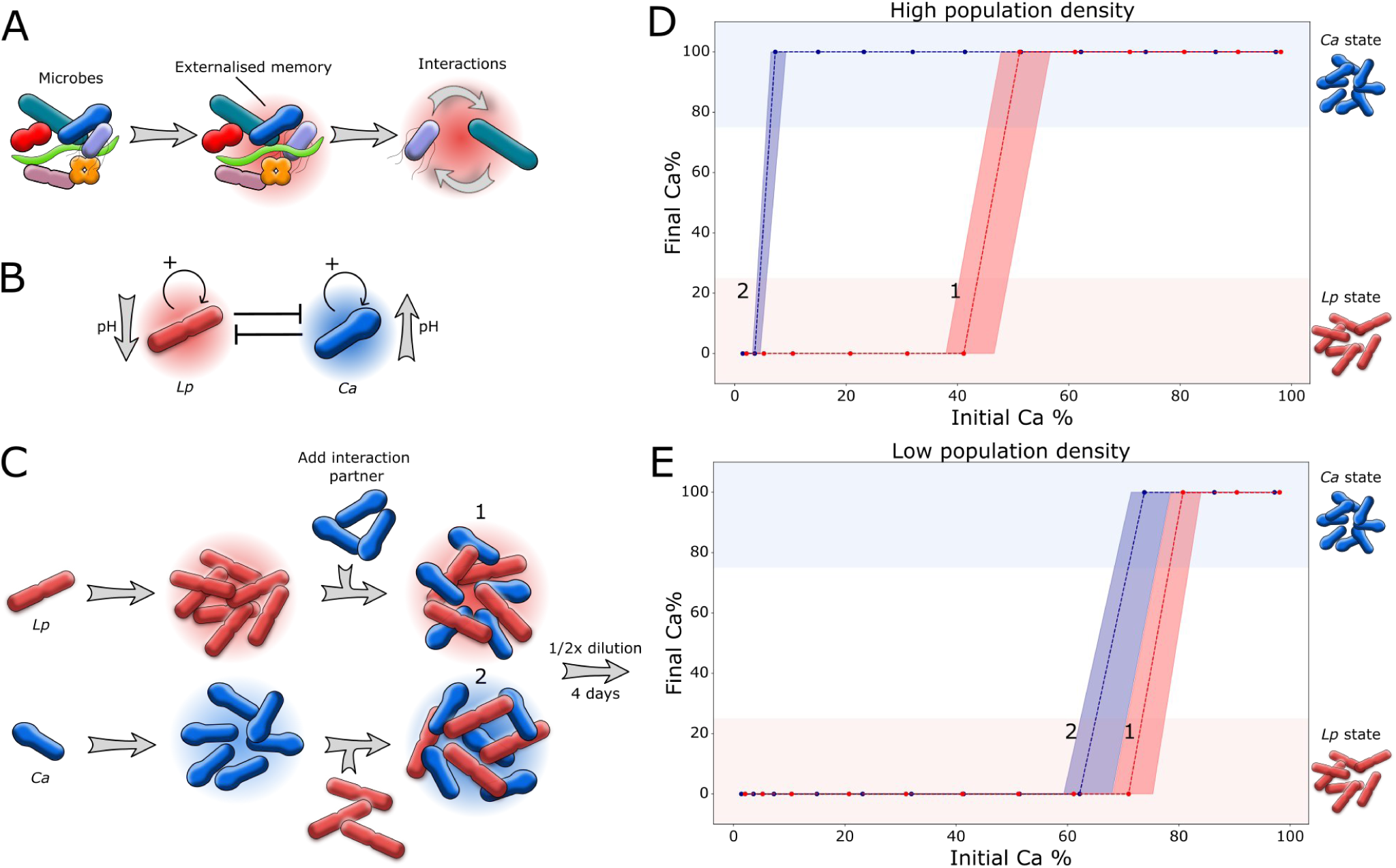
Microbial interaction outcomes depend on the system’s past. **(A)** Microbial communities can chemically change the environment and this chemical modification depends on the microbes’ growth conditions. If this chemical change persists over time, future interactions may be affected by the chemical change and, thus events in the past can impact future interaction outcomes. **(B)** In our two-species model system, *Lp* decreases the pH of the medium, while *Ca* increases pH of the medium, thus inhibiting each other. **(C)** Adding *Ca* to an established culture of *Lp* and *vice versa* can result in different interaction outcomes. When a species interacts in its own memory it has a higher chance to outcompete the interaction partner. However, the effect only occurs at sufficiently high population

We show in the following that such a population memory can indeed originate from relatively simple features of microbial interactions and impact microbial interactions based on past events. Population memory can lead to surprising reactions of a bacterial community to temperature or oxygen. Finally, we could show that population memory can drastically alter the assembly of complex communities.

### Microbial interaction outcomes are shaped by the system’s past

To explore the idea of population memory, we started with a simple model system consisting of the two microbes, *Lactobacillus plantarum (Lp)* and *Corynebacterium ammoniagenes (Ca).* We used this model system in the past to study microbial interactions. In media that contain glucose and urea as major carbon and nitrogen sources (soytone and yeast extract being minor C and N sources, respectively, Methods), *Lp* acidifies the environment by secreting organic acids, and *Ca* alkalizes the media by cleaving urea into ammonia (Fig. 1B). Since *Lp* and *Ca* prefer acidic and alkaline environments, respectively, the two species produce environmental pH values that they can tolerate, but the other species cannot. Accordingly, both species cannot coexist, and one outcompetes the other, where the initial relative abundance determines the winner^20^.

To test whether a different past can change the fate of this 2-species community, we performed the experiment depicted in Fig. 1C (and described in more detail in the Methods). *Lp* and *Ca* were grown independently, free to change their environment. Afterward, cells of the interaction partner were added without their environment (e.g., after washing them) to obtain communities with similar species abundances (Fig. 1B). Despite having similar compositions; the communities differ in how they were built and, thus, their past. The experiment was performed for different fractions of the two interaction partners (see Methods). After mixing the interaction partners, all the communities were treated the same and grown under daily dilution into fresh media for four days. The final community composition was analyzed by counting colony-forming units (CFU, see Methods). The results are shown in Fig. 1D.

First, as an overall trend, we can see that the competition outcome depends on the initial mixing ratio of the two species. The higher the initial abundance of a species, the more likely it wins the competition, e.g., at a high initial abundance of *Lp*, *Lp* tends to win, and at a high initial abundance of *Ca*, *Ca* tends to win. However, the interaction outcome is not only determined by the relative abundances of the two species. In the range from around 4% to 46% initial abundance of *Ca*, the system shows different outcomes depending on the system’s history, even for the same initial relative abundances. The species that previously modified the environment had a higher chance to win, thus exhibiting that the system’s past impacts the interaction outcome.

To further investigate if the collective change of the environment drives this memory effect, we repeated the experiment at lower cell densities. If collective actions of the communities are necessary for memory formation, it should be weakened or disappear at low cell densities. Fig. 1E shows that this is indeed the case. The interaction outcomes are the same, independent of the system’s history (Supplementary Fig. 1). Moreover, the supernatant of one species is sufficient to favor this species in competition, showing that only externalized but no cellular memory is required to explain the observed effect (Supplementary Fig. 3). Thus, we can show here that microbes can build up an externalized memory as a consequence of collective action that impacts future interaction outcomes.

The results of Fig. 1 show how a community assembles can lead to different outcomes even for the same initial composition. Accordingly, the history of a microbial community can decide its future composition. These observations raise the question of whether a similar mechanism can cause microbial communities to develop differently based on past events like different growth conditions. The metabolic state of bacteria changes with environmental conditions such as temperature^21–23^, oxygen concentration, and the availability of resources. Accordingly, how microbes change the environment depends on these growth conditions, and different past growth conditions may impact future interaction outcomes differently. To test this idea, we grew our two model strains, *Lp* and *Ca*, under different conditions each. The temperature was either kept at 18°C or 35°C, with the latter being preferred by both strains. The oxygen levels were either kept low (preferred by *Lp*)^24^ or at atmospheric conditions (preferred by *Ca*) as described in the Methods. *Lp* grown under preferred conditions was mixed with *Ca* grown under unpreferred conditions and *vice versa* (Fig. 2A). In this way, similar community compositions were obtained, but from bacteria grown under different conditions (Fig. 2A). After mixing, the communities were all grown under daily dilution for four days. As shown in Fig. 2B, different past growth conditions result in different interaction outcomes. Again, this memory effect disappears upon dilution, e.g., at low cell densities (Supplementary Fig. 2). To understand better how the different growth conditions could impact the interaction outcomes, we measured how the bacteria change the pH–the major driving factor of bacterial interactions in this system^20^. As shown in Fig. 2C, the pH change per colony-forming unit is higher in preferred vs. unpreferred conditions for both species. These results suggest that past growth conditions–in this case, temperature and oxygen–can lead to different environmental changes by the bacteria and, thus, different future interaction outcomes.

**Fig. 2:**
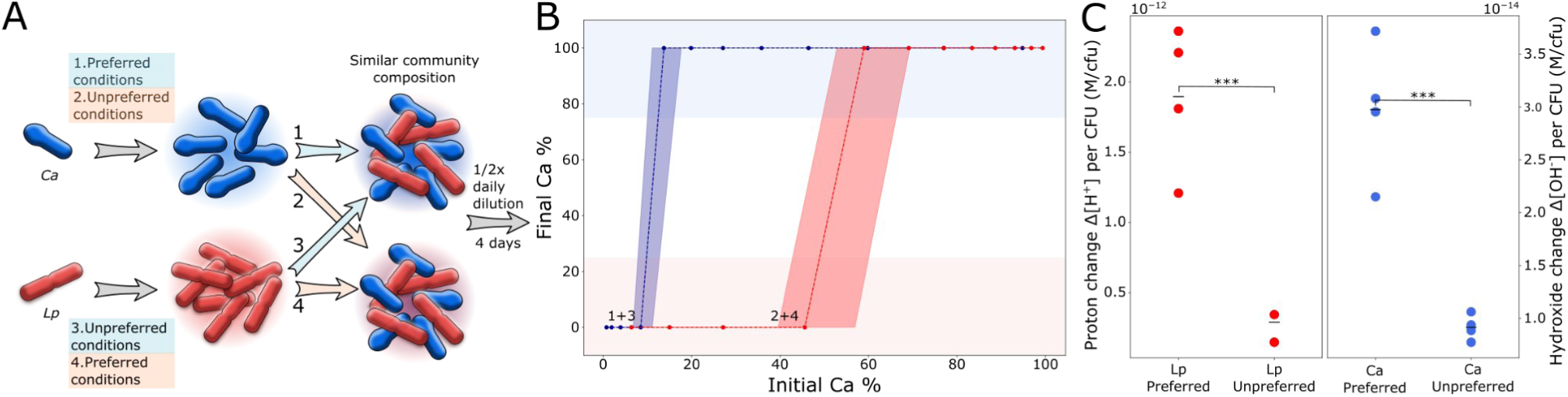
Past growth conditions impact microbial interactions through population memory. **(A)** *Ca* grown in preferred conditions was mixed with *Lp* grown in unpreferred conditions and *vice versa* as described in the methods. **(B)** Population memory formed in preferred conditions enables *Ca* to outcompete *Lp* at lower relative abundances as compared to when *Ca* memory is formed under unpreferred conditions. The effect diminishes with increasing dilution of memory with fresh media (Supplementary Fig. 2). Dashed lines shows again the median curves and shaded area minimal and maximal curves. **(C)** In order to study the effect of different growth conditions on the change of the environment, both bacteria were grown in their corresponding preferred and unpreferred conditions. Preferred conditions were marked by greater proton change in case of *Lp* (Student’s t-test, p-value=0.000865) and greater hydroxide change per CFU in case of *Ca* (Student’s t-test, p-value=0.000747) as compared to that of unpreferred conditions. Data points represent biological replicates.

### A simple mathematical model shows population memory

To get a better mechanistic understanding of the observed memory effect, we formulated a simple mathematical model. In this model, species interact only through the environment, e.g., they change an environmental variable *p,* where the variable’s value determines the bacteria’s growth (Fig. 3A). The bacteria can only grow if this environmental variable lies within a specific range. The corresponding differential equations are shown in Fig. 3A. *N_i_* is the population density of species *i*, *p* can be understood as a metabolite concentration in the environment that is changed by the bacteria and *p_o,i_* is the optimal value of *p* for species *i*. 𝜭 is the Heaviside function that becomes equal to one only if its argument is positive, that means if *p* is close enough to *p_o,i_*. *r_dilution_*corresponds to a dilution factor that removes cells from the system and, therefore, equals the death rate of the bacteria. At the same rate *r_dilution_*, fresh media with *p*=0 (note that negative *p* is allowed in this model) is added to the system. The environmental parameter *p* is changed by the species *i* with the rate *k_i_*.

**Fig. 3:**
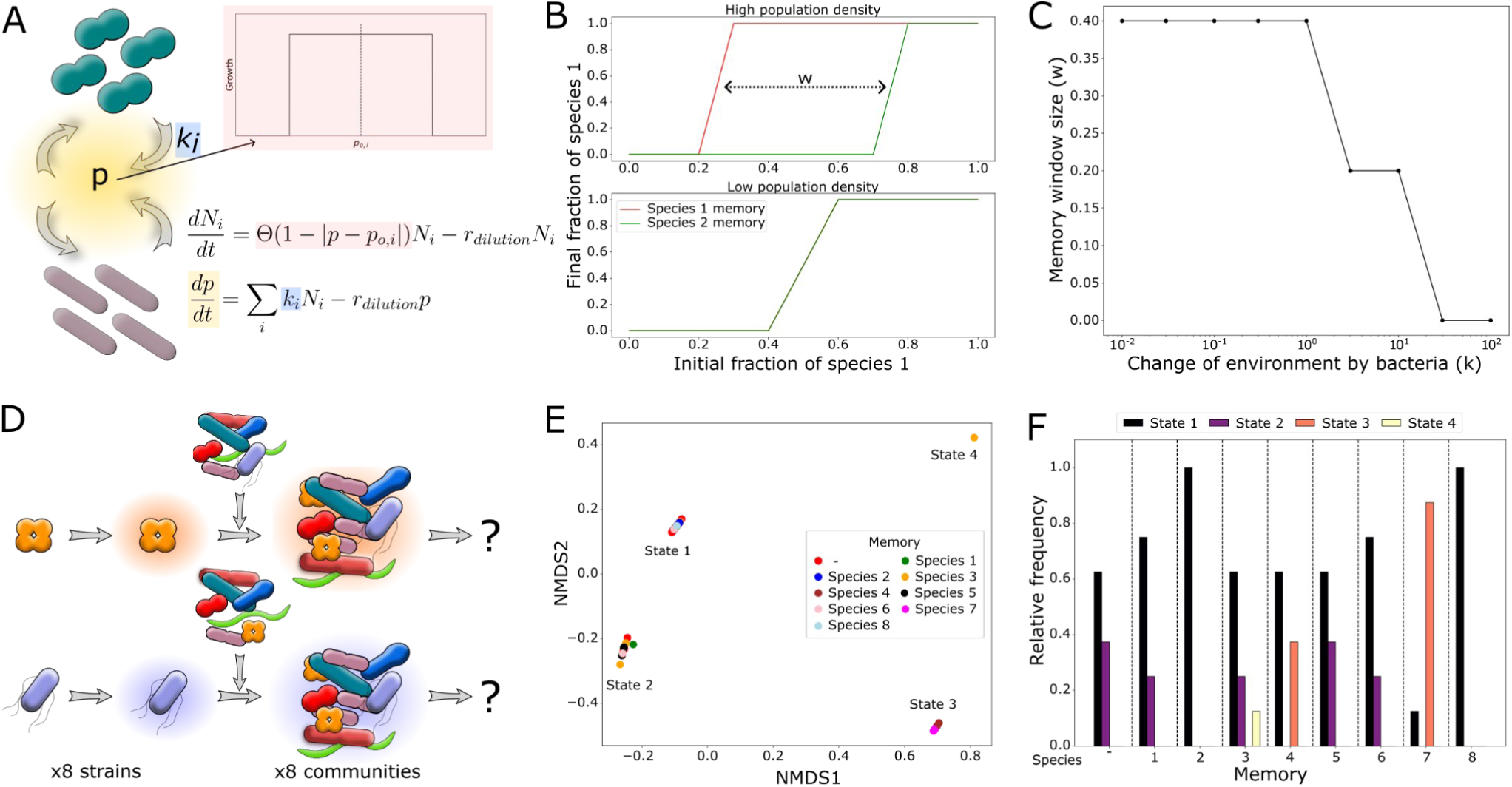
A simple model can recapitulate memory effect in interactions and suggests that it affects community assembly. **(A**) In order to simulate microbial interactions in the presence of collective memory, the shown equations were used, where the growth rate of species N*_i_* is determined using the Heaviside step function, 𝜭, of the difference between metabolite concentration *p* and optimal metabolite concentration *p_o,i_.* The rate of change in metabolite concentration is governed by *k_i_* and dilution rate *r_dilution_*. This system ensures maximum growth rate of the species when the concentration of the metabolite *p* is close to the respective optimum value for the species, as illustrated in the *p* vs growth plot. **(B)** Simulating the equations for two mutually-excluding species and one metabolite for different initial mixing ratios reveals that a species has a higher chance to outcompete the other species in the presence of its own memory, recapitulating the memory effect observed experimentally. **(C)** A faster change of the environment by the bacteria *k* (*k_1_* = *k*, *k_2_* = *-k*) removes the memory effect. The window size shown on the y-axis corresponds to the difference between the two curves for both species’ memory as depicted in (B). **(D)** Extending the complexity to eight species and three environmental metabolites, we simulated the assembly of several multispecies communities

With this simple model, we first tried to recapitulate our experimental findings. Accordingly, we simulated the interaction of two species that lower or increase *p* and also prefer low or high *p* v alues (*p_o,i_*), respectively. The two species, therefore, change the environment in a way that is beneficial for themselves but detrimental for the interaction partner. We first grew one of the species and mixed it with cells of the other species as done experimentally before (Fig. 1, Methods). Indeed, we observed a similar memory effect in the model that also disappears upon diluting the system (Fig. 3B). It is important to note that such a memory effect where for the same initial species abundances, different interaction outcomes are obtained is only possible because the interaction is mediated through the environment. The environment acts as a third variable (Supplement). Moreover, the memory effect only occurs if the change of the environment by the species (*k_i_*) is not too fast (Fig. 3C). If the bacteria change the environment too rapidly, the memory is “overwritten” by the bacteria and thus cannot influence the interaction outcomes (see also Supplement).

Having this model at hand prompted us to investigate the significance of such a memory effect for the assembly of more complex communities. Accordingly, we repeated the simulations for eight species systems. One species was grown, and then cells of the other species were added to the system to obtain the same initial abundances but with different histories (Fig. 3D). This was repeated for different abundances of the same set of species (Supplementary Fig. 7A). The dynamics of the system were simulated to obtain the final composition at the end of the simulation run (Supplementary Fig. 5). These final states are visualized in Fig. 3C after reducing the 8-dimensional outcome (one dimension for each species) to a 2D plot by applying Non-Metric Multidimensional Scaling (NMDS) to the Bray-Curtis dissimilarity matrix of the obtained final communities (Methods).

We first notice that this model system shows multistability even without memory. When the cells are mixed in different ratios in fresh media, the system ends up in one out of two final states (Fig. 3E red and Supplementary Fig. 7F). The frequency for the different initial community mixtures to end up in one of the states is given in Fig. 3F, left. In the presence of population memory, we observe two interesting phenomena. First, the relative frequency of ending up in one of the two alternative states changes depending on which species’ memory the assembly takes place in. Second, the systems can end up in novel states that could not be reached in the absence of memory (yellow and red bars in Fig. 3F). Similar findings were observed in simulations with a second set of microbes (Supplementary Fig. 8). Overall, these simulations suggest that the assembly of the same set of species with the same initial abundances can nevertheless end up in different communities in the presence of population memory.

### Population memory can impact community assembly

To test if population memory can indeed affect the assembly of more complex microbial communities and to study further how common this phenomenon is in an arbitrary set of microbes, we performed a lab experiment that equals the simulation shown in Fig. 3. We randomly selected eight bacterial species from our *C. elegans* gut strain collection for community assembly. To estimate the impact of population memory on the assembly process, the eight species were mixed in different abundances, where the memory of only a single species was present in the system during assembly, equivalent to Fig. 3D. This corresponds to a situation where one species grows first in its habitat and changes the chemical composition, followed by the arrival of the other bacteria. We tested six initial community compositions (Supplementary Fig. 13A), with eight different population memories in 3 replicates each. This results in 144 community assembly processes, along with the replicates assembled in the absence of memory. After mixing we cultivated the communities for 12 days under daily dilution into fresh media, e.g., all communities were treated the same after the initial setup. The compositions of the communities were assayed on days 5, 10, and 12 of the experiment by plating on NM agar and counting the forming colonies (Methods, Supplementary Fig. 9). Bray-Curtis dissimilarities between the communities of day 12 were calculated, followed by dimensionality reduction using NMDS (Fig. 4A). The data for the remaining days and all raw data are shown in Supplementary Fig. 15 and Supplementary Figs. 11-13. As can be seen, the communities converge towards two final states, e.g., the system is multistable. The two states are differentially colored in Fig. 4A based on the hierarchical clustering of the final communities’ Bray-Curtis dissimilarities (Supplementary Fig. 14C, Supplementary Fig. 10). This observation also aligns with the visual impression of the raw data (Supplementary Fig. 13). Multistability seems to be a relatively common feature of microbial communities, as we will discuss in a separate publication. The different initial communities collapse either into state 1, which is dominated by *Rouxiella badensis (Rb)*, or state 2, which is dominated by *Arthrobacter nitroguajacolicus (An), Rhodococcus qingshengii (Rq)* and *Sphingobacterium multivorum (Sm)* (Fig. 4B). In the absence of memory, the communities end up mostly in state 1 and only in around 22% of the cases in state 2 (Fig. 4A, Fig. 4C left). The states in which the communities end up are mainly determined by the initial species composition (Supplementary Fig. 14E).

**Fig. 4:**
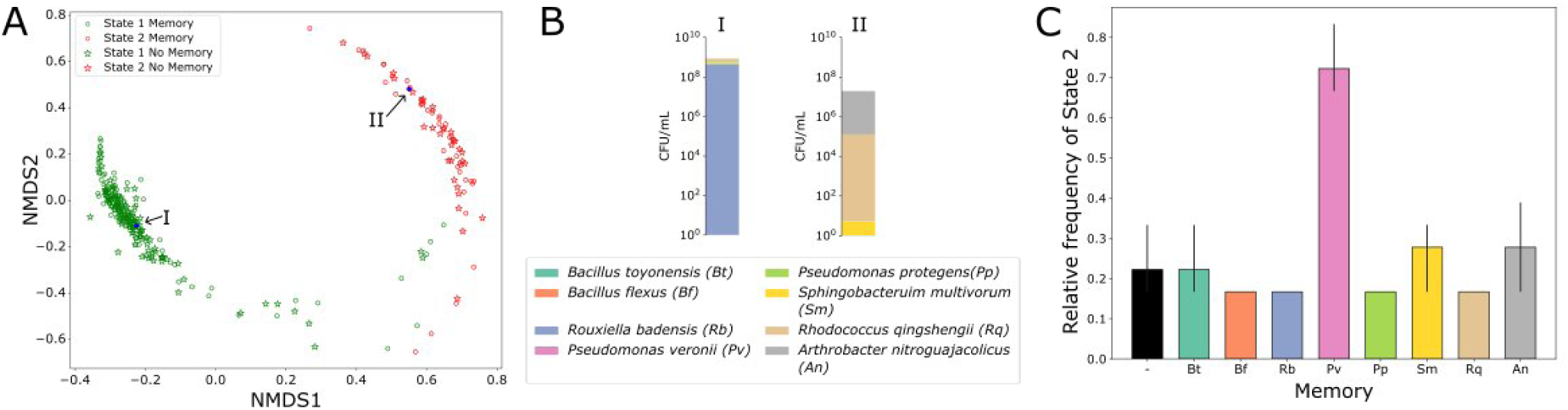
Collective memory impacts community assembly. **(A)** Eight-species communities were mixed with varying initial compositions either in the absence or presence of different population memories. NMDS plots based on Bray-Curtis dissimilarities between final community compositions depict the communities ending up in two final states, referred to as state 1 and state 2. The state for each community is obtained by hierarchical clustering performed on Bray-Curtis dissimilarities (Methods). **(B)** State 1 is dominated by *Rouxiella badensis (Rb)* with relatively high total cell densities, and state 2 is dominated by *R. qingshengii (Rq)*, *S. multivorum (Sm)*, *A. nitroguajacolicus (An)* with relatively lower total cell densities. **(C)** Exposure to the population memory of *P. veronii* during the initial community assembly increases the frequency of community ending up in state 2 as compared to absence of memory. Lines on bars represent

In the presence of memory, however, the outcome changes (Fig. 4C, Supplementary Fig. 14D). In particular, the memory of *P. veronii* significantly increases the frequency of communities ending up in state 2 (ANOSIM^25^, calculated on Bray-Curtis dissimilarities, p-value = 0.001, R-value = 0.4261). In contrast, communities assembled under other memories were not significantly different from those assembled without memory. Overall, the population memory of specific strains can strongly impact community assembly, whereas, at the same time, different communities can also be variably susceptible to this memory effect (Supplementary Fig. 13). Although the memory of *P. veronii* impacts the community assembly, *P.veronii’s* memory does not ensure its survival in the final community (Supplementary Fig. 13).

## Discussion

Despite their simplicity, bacteria can store memory in many ways^26,27^, allowing them to retain cellular states over long periods of time^6,8,28,29^. Bacteria’s growth and gene expression can depend on their past growth conditions^30–33^, and the virulence of pathogenic bacteria can be lastingly altered by a previous host contact^34^. Bacteria can store these memories inside the cell, for example, by retaining active proteins in the cytosol over longer times or showing persistent gene activation even after the stimulus dissapeared^31,35,36^. In many cases, however, the mechanistic origin of the memory is not understood^30,34,37^.

This work describes a very different type of memory that is not stored inside the cell but externalized in the environment. Bacteria can generate external memory by collectively changing the chemical composition of the environment^13,38–40^. These environmental changes may depend on events in the past that, in this way, impact interaction outcomes in the future. Since single bacteria can usually not significantly change the environment, whole populations have to work together to form this external memory, which we call population memory accordingly. Microbes are well known for collective traits^41^, like cooperation^42^ or quorum sensing^43^, which aligns with their community lifestyle^44^. Our findings add another collective behavior - e.g., population memory - to this list.

Related to our population memory, a “collective memory” has been theoretically described for animal groups such as fish swarms. In such animal groups, specific swarming patterns may appear and persist over time only due to the interactions amongst the groups’ individuals but without the memory stored inside the individuals^45^. Another example of externalized, collective memory in nature is pheromone trails, which occur in social insects^12^. Thus, ants can deploy a pheromone trail back to their nest upon finding food to guide other ants, although these ants do not have an intrinsic memory of the food source. The volatility of pheromones sets the period of this collective memory; the faster they evaporate, the shorter the memory^9^. Pheromones are particular substances that impact behavior. We, however, show that even rather unspecific by-products of metabolism can function as ingredients of population memory in microbes.

Population memory is not just an exciting concept; it may have far-reaching ecological consequences, particularly concerning microbial interactions and community assembly. Thus, the ability of a new invader to establish itself in a community may not just depend on the community’s composition but also its past. In that regard, population memory can be regarded as an extension of the priority effect^47–49^. In priority effect, the order of arriving species determines the assembly of communities. In contrast, in population memory, essentially, any past event or growth condition that impacts microbial metabolism can alter community assembly.

Ecosystems are always the outcome of their historical development; for example, in macroscopic ecosystems, past rainfalls^50^ or land usage^51^ can impact future ecosystem development. The presence or absence of species in an ecosystem also depends on whether these species historically invaded this ecosystem or died out at some point^52^. Population memory, e.g., an externalized and collective memory produced by an ecosystem’s organisms, adds another mechanism that could shape trajectories even in macroscopic ecosystems. But, outside our very controlled lab experiments, it may often be difficult to distinguish if a past event impacted community assembly directly, e.g., by impacting bacterial growth or indirectly via the formation of population memory.

One can also understand our results in the context of niche construction. Niche construction describes the alteration of the environment by organisms^53,54^. Microbes that change the chemical composition of their environment construct an ecological niche^13,20,55,56^. We show here that different niches could be constructed depending on the past conditions, and differently constructed niches accordingly impact the future development of ecosystems differently. Similar statements have been made about human niche construction, as in the case of agriculture, where human modifications of the environment can be a response to changing environmental conditions (e.g., climate change), and anthropogenically altered environments can impact humans and nature over a long time^57,58^. Therefore, we may regard niche construction less as resulting from fixed traits of organisms but as a more flexible process that connects the past conditions of an ecosystem with its future.

On the theoretical side, population memory may have overlooked consequences. Often, Lotka-Volterra models are used to model microbial communities for theoretical insights^59^ or to predict experimental outcomes^60–62^. However, these models only account for direct species interaction–e.g., not mediated through the environment–and thus cannot describe population memory. Indeed, Lotka-Volterra models can strongly deviate from models that consider environmental changes because they ignore lasting environmental changes ^63,64^.

The microbial communities in our guts are connected to our health in many ways^65,66^, and accordingly, long-lasting changes within this microbiota could impact our well-being^67^. Indeed, the gut microbiota has been shown to possess a memory of past nutrient exposure, medical treatments, or infections^68–71^. Still, given the intrinsic complexity of these systems, it is very challenging to identify the underlying mechanisms. At least in a simple model system of a bacterial gut community, memory was suggested to be caused by the interplay between bacterial metabolism and modification of the environment^69^. Population memory may, therefore, also play a role in host-associated microbial communities and be of medical relevance. In particular, medical treatments or dietary changes^72,73^ can alter the pH in the gut microbiota, which can cause population memory, as we showed here. Further studies are required to explore the potential role of population memory in the resistance against pathogen invasions or the success of medical procedures such as fecal transplants.

## Materials and Methods

### Media and buffers

Nutrient media (NM) was prepared with 10g/L Yeast extract (J23547, Thermo Scientific, Kandel, Germany) and 10g/L Peptone (91079-46-8, Merck Millipore, Darmstadt, Germany) mixed in distilled water and pH adjusted to 7. NM was then autoclaved at 121℃ for 20 mins at 103kPa. Base media (BM) was prepared by mixing 1g/L Yeast extract, 1g/L Soytone, and 10mM NaH_2_PO_4_ (28011.291, VWR, Darmstadt, Germany) in distilled water, and the pH was adjusted to 6.5 (8172BNWP, 2115001, Thermo Scientific, Braunschweig, Germany). BM was autoclaved as described for NM above. Before usage, 0.1mM CaCl_2_ (A4689.0250, PanReac AppliChem, Darmstadt, Germany), 2mM MgSO_4_ (11596, Thermo Scientific, Kandel, Germany), 4mg/L NiSO_4_ and 50mg/L MnCl_2_ were added.

Supplemented Base media (SBM) was prepared by adding 1% w/v D-Glucose (0188-2.5KG, VWR, Darmstadt, Germany) and 0.8% w/v Urea (U/0500/53, Fisher Scientific AG, Reinach, Switzerland) to Base media and vacuum filtering through 0.2µm filtration unit (Filtropur V50 83.3941.001, Sarstedt, Nümbrecht, Germany).

NM Agar plates were prepared by dissolving 10g/L Yeast extract, 10g/L Peptone, 10mM NaH_2_PO _4_, and 2.5% Agar (5210.2 Carl Roth, Karlsruhe, Germany) in 980mL water. The mixture was autoclaved, and 50mL solution was poured into each 150x200mm petri dish (82.1184.500 Sarstedt, Nümbrecht, Germany). Before usage, plates were dried at room temperature for 3 days. For pH-selective NM Agar plates, the pH of the mixture was adjusted to either 5 or 10 before autoclaving, and after autoclaving, 0.02% sterile Glucose was added to the mixture upon cooling.

To make M9-K minimal media, 43mM NaCl (12314, Alfa Aesar, Thermo Scientific, Kandel, Germany), 54mM KCl (1.04936.1000, Supelco, Merck SA, Darmstadt, Germany), 93mM NH _4_Cl (21236.267, VWR, Darmstadt, Germany), and 10mM NaH_2_PO_4_ were dissolved in distilled water, pH was adjusted to 7, filled up to 1L with distilled water, and autoclaved as described for NM media above. After cooling down, the sterile addition of 1mL/L 1000x trace metals mix (Teknova, Hollister, California, USA T1001), 2mM MgSO_4,_ and 0.1mM CaCl_2_ was carried out.

Tryptic soy broth (TSB) was prepared by mixing 17g/L Tryptone (95039-1KG-F Sigma Aldrich, Merck SA, Darmstadt, Germany), 3g/L Peptone, 5g/L NaCl, 2.5g/L K_2_HPO_4_ (191431, MP Biomedicals, Eschwege, Germany), and 2.5g/L D-Glucose in 1L distilled water. The medium was sterilized using vacuum filtration through a 0.2µm filtration unit.

Phosphate Buffered Saline (PBS) + 0.4% Tween-80 was prepared by diluting 8g/L NaCl, 200mg/L KCl, 10mM Na_2_HPO_4_ (30435-1KG, Honeywell, Seelze, Germany), 1.8mM KH_2_PO_4_ (26923.298, VWR, Darmstadt, Germany) and 0.4% v/v Tween-80 (8.22187.0500 Sigma-Aldrich, Merck SA, Darmstadt, Germany) in distilled water, followed by sterilization with autoclave as described for NM above.

### Bacterial strains

*Lactobacillus plantarum* (referred to as *Lp*) is a gram-positive, non-motile, lactic acid-producing bacterium acquired from the American Type Culture Collection (ATCC8014). *Corynebacterium ammoniagenes* (referred to as *Ca*) is a gram-positive aerobic bacterium and was also obtained from the American Type Culture Collection (ATCC 6871).

Gut strains were isolated from *C. elegans,* as described by Ortiz et al.^27^. Eight strains (*Bt, Bf, Rb, Pv, Pp, Sm, Rg, An*) were selected to ensure different colony sizes, colors, and morphology - to distinguish them upon colony counting as described below, as well as ensuring phylogenetic diversity (Supplementary Fig. 9, 13B). Bacteria were streaked on NM Agar plates, and colonies were allowed to form by incubation at 30°C for 24 hours. The plates were subsequently stored at 4°C for up to 1 week. Single colonies from these plates were used to start the experiments described below.

### Two-species memory assay

Single colonies of *Lp* and *Ca* were grown in 5mL NM overnight in 50mL tubes (62.547.254, Sarstedt, Nümbrecht, Germany) at 30°C shaking at 225rpm (Innova 2000, New Brunswick Scientific, Nürtingen, Germany). The lids were left open for 1/8th of the turn and fixed with tape. After 18 hrs, bacterial cultures were centrifuged at 2451 rcf for 3 mins and resuspended in 5mL SBM. Resuspended cultures were 1/100x diluted into 2 x 5mL fresh SBM cultures for each species in 50mL tubes for incubation at 30°C on shaking for 24 hours. The following day, all 5mL cultures from the same species were pooled and split into two tubes - ‘washed’ cultures and ‘unwashed’ cultures.

To make ‘washed’ cultures, bacterial cultures were centrifuged at 2451 rcf for 3 mins (Centrifuge 5810, Eppendorf, Hamburg, Germany), and the supernatant was taken out and filtered into fresh 50mL tubes using 0.2µm syringe filter (83.1826.001, Sarstedt, Nümbrecht, Germany). Pelleted bacteria were resuspended and washed twice with fresh SBM. OD_600_ was taken using Implen OD_600_ reader for washed and unwashed bacterial cultures. OD_600_=1 cultures were prepared for washed samples using fresh SBM and with respective supernatants for unwashed samples for each species. Dilution rows for OD_600_=1 cultures were prepared with PBS + 0.4% Tween and plated with 5μL droplets using Viaflo 96 for initial CFU counts on pH 5 NM plates for *Lp* and pH 10 NM plates for *Ca*. *Lp* “unwashed” was mixed with *Ca* “washed” culture and vice versa in increasing percentages from 2% to 98%. Mixed cultures were diluted with SBM with ratios of 1/2x, 1/6x, 1/20x, and 1/100x in 96 deep well plates (786201, Greiner Bio-One Frickenhausen, Germany). The plate sealed with breathable sterile rayon film (VWR 391-1262) was incubated on shaking at 1250rpm on a platform shaker (Titramax 100, Heidolph Instruments GmbH, Schwabach, Germany) at 30℃. 1/2x dilutions with fresh SBM were carried out daily for 4 days. Dilution plates were made with 1/10x dilution in PBS + 0.4% Tween in 96 well microtest plates (82.1581.001, Sarstedt, Nümbrecht, Germany), followed by droplet plating of 96 well plates on pH selective NM Agar plates with 5uL droplets using Viaflo 96. pH selective NM Agar plates were incubated at 30℃ for 48 hours before CFU counting using a stereo microscope (TL3000 Ergo, Leica Microsystems, Wetzlar, Germany). From observed CFU/mL values, actual *Lp/Ca* ratios, and percentages were calculated.

### Two species memory assay under different growth conditions

For studying the effect of temperature on population memory, two SBM cultures were prepared for each species (after primary culture in NM, as described above) – one grown in preferable conditions and the other grown in unpreferable conditions, as described in Fig. 2A. Depending on the different physiological requirements of the bacteria, 2 x 50mL tubes (per condition per species) were either covered with breathable rayon film (Ca_Pref_ and Lp_Unpref_) or completely closed (Lp_Pref_ and Ca_Unpref_) during incubation at either 35℃ (Ca_Pref_ and Lp_Pref_) or at 18℃ (Ca_Unpref_ and Lp_Unpref_) on shaking at 225rpm for 24hrs.

The following day, bacterial cultures of the same species and conditions were pooled and centrifuged at 2451 rcf for 3 mins, and half of the supernatant per tube was taken out and filtered into fresh 50mL tubes using syringe filters. Optical density reading was taken, and using their respective filtered supernatants, OD_600_ = 1 was adjusted for both species at both conditions, and dilution plating for initial CFU counts was performed as described above. Ca_Pref_ was mixed with Lp_Unpref_, and Ca_Unpref_ was mixed with Lp_Pref_ in increasing percentages from 2% to 98%, and mixed cultures were diluted with SBM with ratios: 1/2x, 1/6x, 1/20x, 1/100x in 96 deep well plates. The following steps were similar to those outlined above.

### Mathematical model, parameter selection, and simulation

Simulations were performed in Python 3.9.7 based on the mathematical model described in Fig. 3A.

To recapitulate the 2-species system, the number of interacting species *N_i_* was set to i = 2. Both species influence one environmental metabolite *p*. *k* and *p_o_* were set to [-1, 1] and [-3, 3], corresponding to two species that change the environmental variable p in opposite directions (as in Fig. 1B) at an equal absolute rate k_1/2_. Species were grown individually with *p_initial* = 0, *N _i__initial* = 1, and *r_dilution_*= 0.1, and steady-state mean *p_stst_*values were recorded to be utilized as *p_initial* in the presence of population memory in the following. To simulate species interactions, the initial *N_1_* values were set to {0.001, 0.01, 0.1, 0.2, … 0.999}, and correspondingly, initial *N_2_* values were set to 1-*N_1_* (see x-axis of Fig. 3B). The initial *N_i_*, as well as *p_initial*, were subjected to various dilutions with dilution factors of {1, 0.3}, as plotted in Fig. 3B. To obtain the population densities and environmental variable over time the differential equations of Fig. 3A were integrated with solve_ivp module in python. The obtained steady-state values are plotted in Fig. 3B. In an effort to ascertain the role of *k_i_* in the memory window size, simulations were performed over multiple values of *k* ∈ {0.01, 0.03, 0.1, 0.3, 1, 3, 10, 30, 100} such that values of *k_1/2_* are [*-k, k*] for two species in one simulation run (Fig. 3C).

To theoretically explore the assembly of more complex communities, we simulated eight species influencing three environmental metabolites. The values of *k_i_* and *p_o,i_* were chosen from uniform distributions using the numpy random.rand function, with *p_o,i_* values ranging from −2 to +3 and *k_i_* values ranging from −10^-3^ to 10^-3^. To obtain each species’ population memory, single species growth was simulated with *p*_initial = [0,0,0], *N*_initial = 10, *r_dilution_* = 0.1, and their respective *k_i_*and *p_o,i_*values, and steady-state *p_stst_*values were acquired. Then eight different communities were assembled with different initial abundances of the same eight species (with distinctive *k_i_* and *p_o,i_* values) in the memory of a single species. The different initial abundances *N*_initial for each species in the communities were randomly selected from {0.1,1,10} and are shown in Supplementary Fig. 7A and 8A. *p_initial* was set to *p_stst_*values from single species growth. In the absence of memory, *p_initial* was set to [0,0,0]. The python function solve_ivp was utilized to perform simulations on equations in Fig. 3A. The final *N_i_*values were used to calculate Bray-Curtis dissimilarities, and based on that, NMDS was performed using sklearn.manifold.MDS module. The outcomes were plotted using matplotlib module.

### Community assembly and determination of final community composition

For each of the eight *C.elegans* gut strains (see above and Supplementary Fig. 9), precultures (5mL) were grown in 50mL tubes in Tryptic soy broth (TSB) overnight at 30°C on shaking at 225rpm. The following day, TSB cultures were washed in M9-K + 0.05% Glucose and 1/100x inoculated into TSB as well as M9-K + 0.05% Glucose cultures for each species in volumes of 5mL at 30°C for 24hrs at shaking. The next day, M9-K + 0.05% Glucose cultures were centrifuged, and the supernatant was filtered using syringe filters.

Communities with different initial abundances (Supplementary Fig. 14A) were mixed. After shaking at 1800rpm for 2mins, 6.67µL of mixed community cultures was added to respective wells with 93.3µL of respective filtered supernatants and 100µL of fresh M9+0.05% Glucose per well in a 96-deep well plate and sealed with breathable sterile rayon film. Daily dilution of 1/30x was performed into fresh M9+0.05% Glucose for 12 days, and final communities were plated on NM agar plates. NM agar plates were incubated at 30℃ for 48 hrs before CFU identification and counting. To ensure the stability of the final states, plating on NM agar plates was also performed on day 5 and day 10 (Supplementary Fig. 11, 12).

### Data analysis

All the plots were plotted using the matplotlib module in Python 3.9.7, while the illustrations were created on Inkscape. Non-metric multidimensional scaling, hierarchical clustering, and K-means clustering were performed using the scipy module in Python 3.9.7, while ANOSIM was performed using the scikit-bio module in Python 3.9.

## Supplement

**Supplementary Fig. 1:**
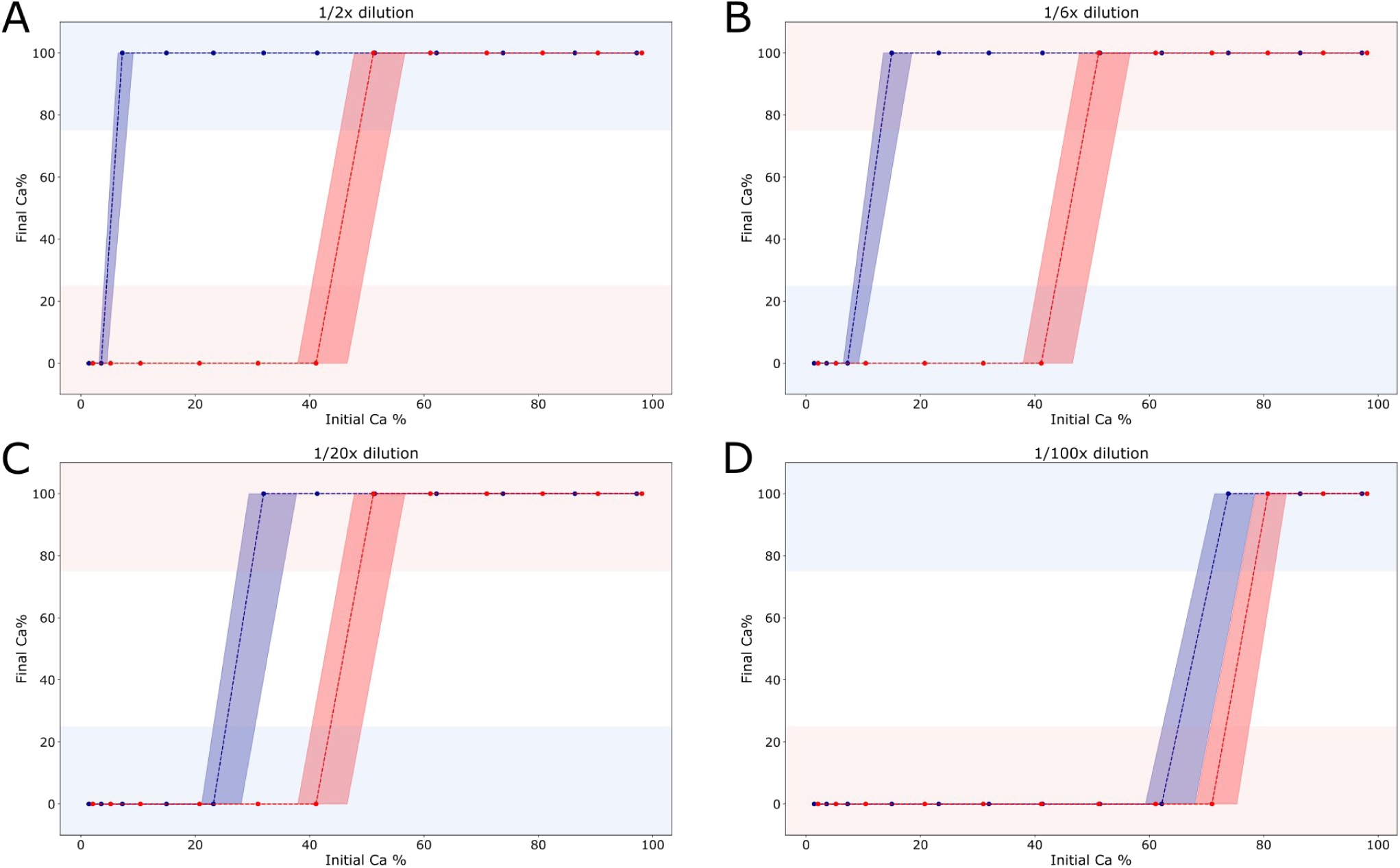
Population memory window decreases from **(A)** 1/2x dilution of memory with fresh media (referred to as High population density in Fig. 1D) to **(B)** 1/6x dilution, **(C)** 1/20x dilution and disappears at **(D)** 1/100x dilution (referred to as Low population density in Fig 1E) reiterating the density dependent role of collective memory in deciding the outcome of microbial interactions.

**Supplementary Fig. 2:**
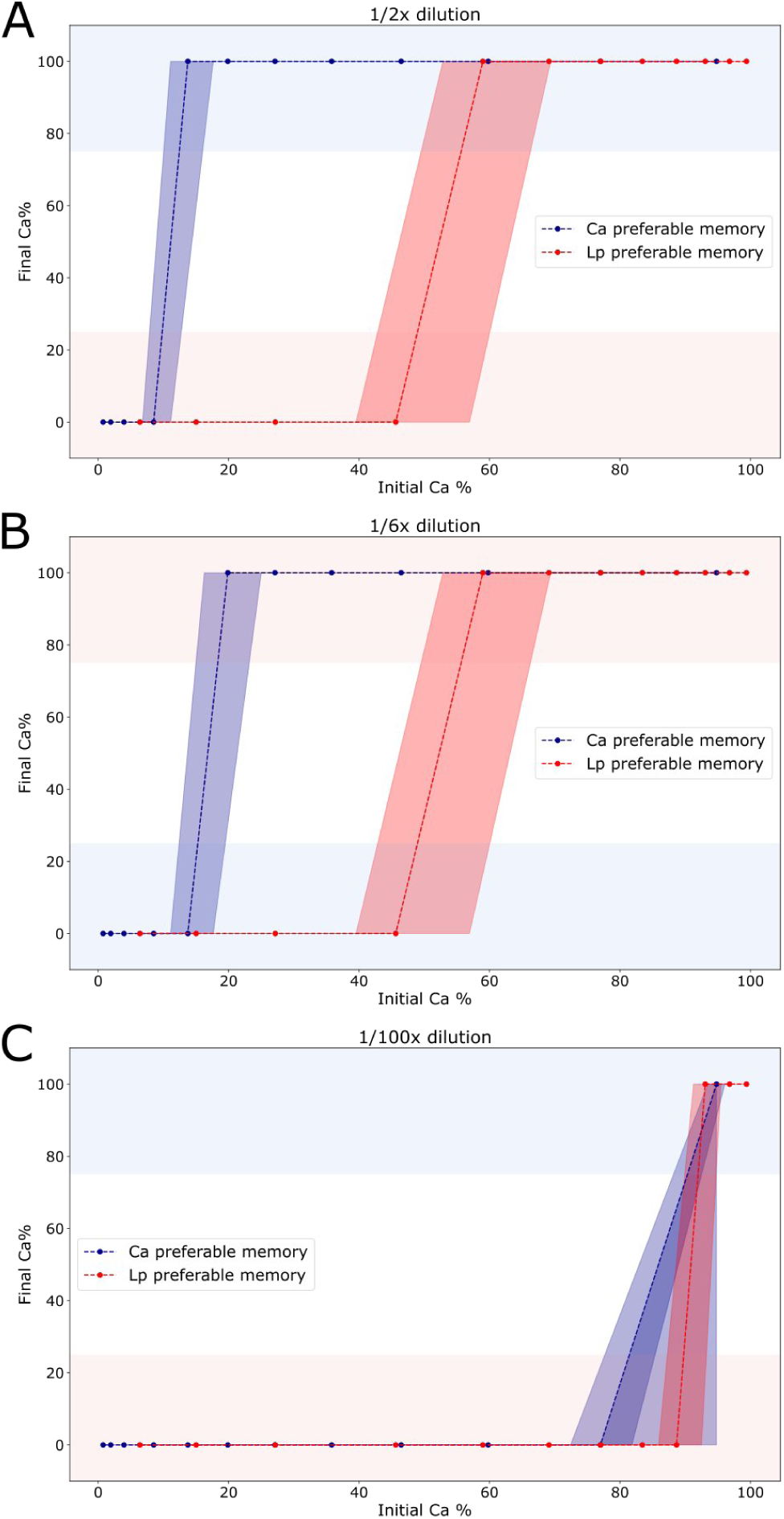
Population memory window decreases from **(A)** 1/2x dilution of mixed memory with fresh media (referred to as High population density in Fig 2C) to **(B)** 1/6x dilution, and disappears at **(C)** 1/100x dilution. Interestingly, memories from both the microbial populations, *Lp* and *Ca,* grown in either preferred or unpreferred conditions are present during microbial interactions.

**Supplementary Fig. 3:**
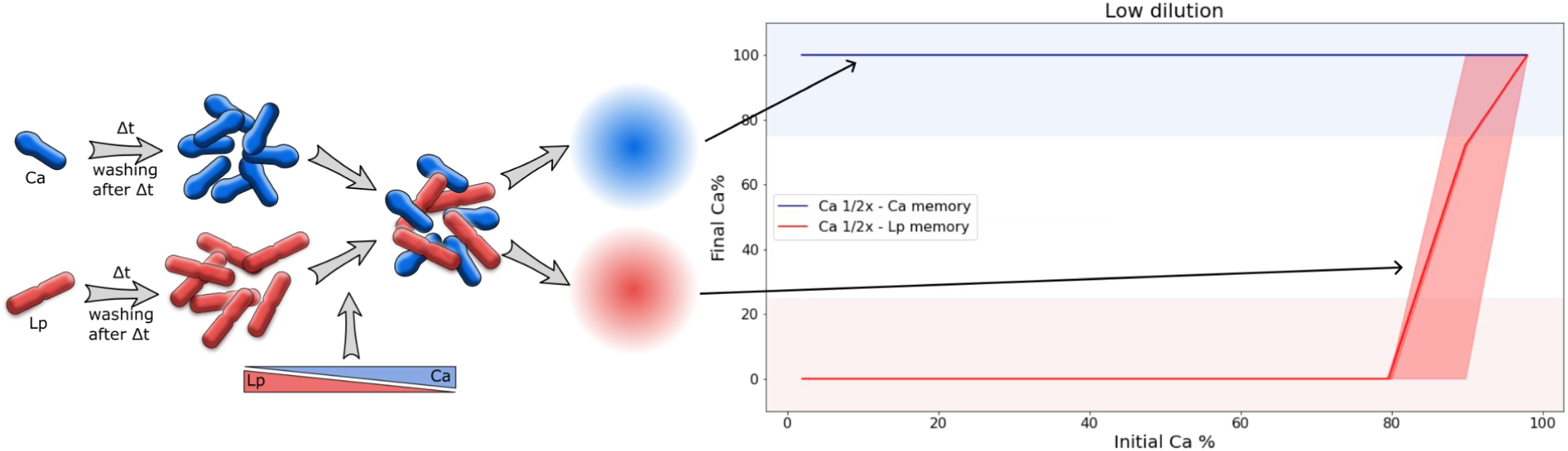
Modified media is sufficient for population memory effect. In order to establish the source of memory effect being externalised memory, we recapitulated experiment in Fig. 1 with different approach. Memories were removed from microbes and added later to the different 2-species communities, thereby establishing similar cellular environment, but different memories. We reestablished the memory effect, where presence of own memory facilitates the microbe in achieving monospecific state.

**Supplementary Fig 4.**
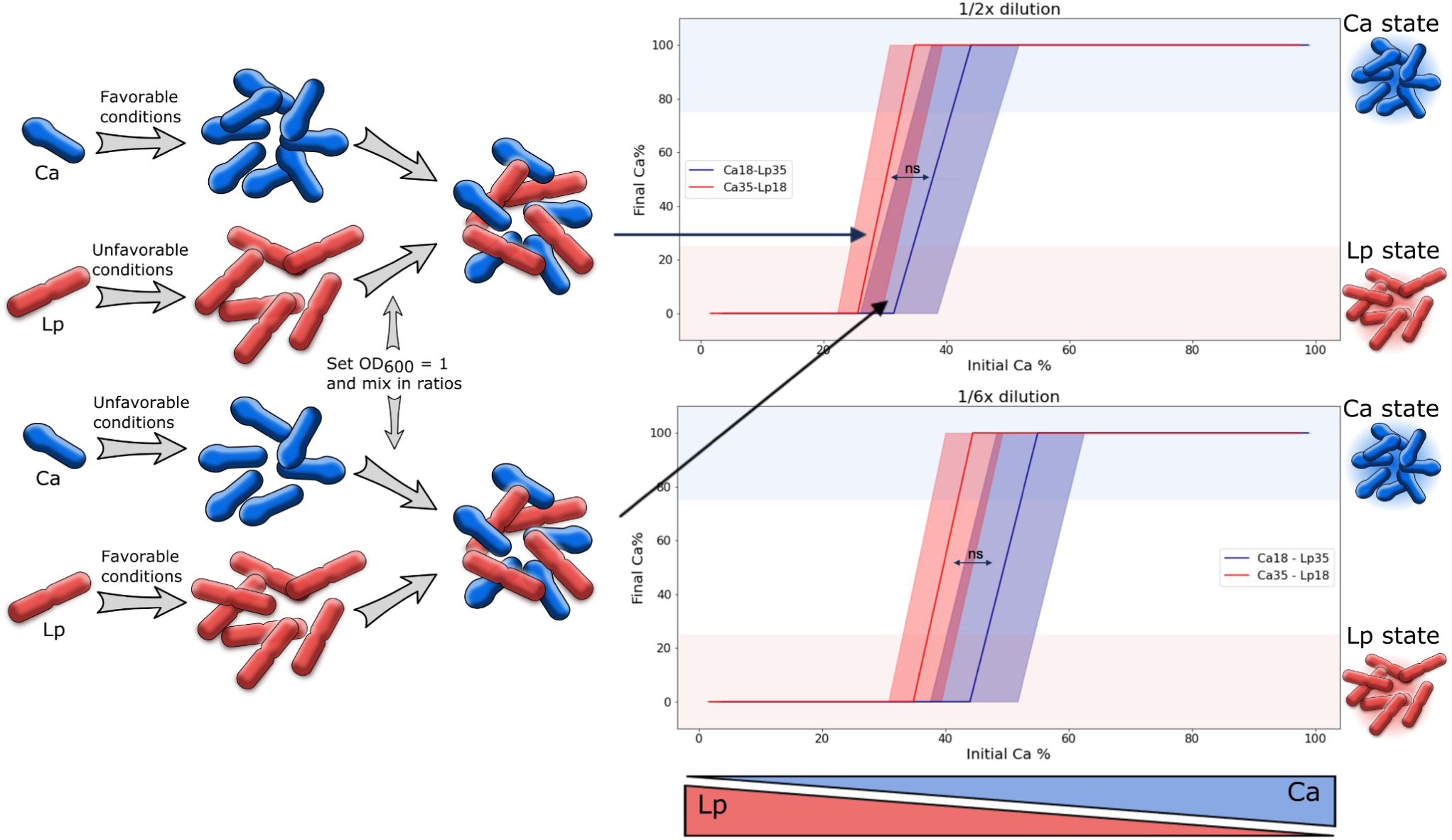
Removing spent media removes memory effect. Aimed at verifying the role of ‘supernatant’ in the memory effect we established an alternative ‘depletion’ approach, where memories were not re added to 2-member communities assembled from microbes grown at favorable and unfavorable conditions. Results demonstrated no memory effect originating from microbes grown in either favorable or unfavorable condition, even at higher cell density.

### Theoretical conditions for the appearance of population memory

For the appearance of the memory effect, two conditions have to be fulfilled:

#### 1) Interactions have to be environmentally mediated

Population memory can lead to different interaction outcomes for the same mixing ratio of a set of N species. Such a memory cannot occur for direct interactions that are not mediated through the environment. In the case of direct interactions the interaction outcomes of N species only depend on the abundances of these species themselves. Therefore for given initial abundances of the N species the same interaction outcomes will be achieved (ignoring noise), because the system’s dynamics only depend on those N variables and no other variables are present.

This is for example true for a Lotka-Volterra type system:

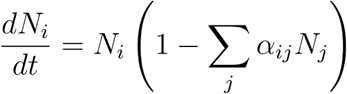

Where the dynamics of the species *N_i_* only depend on the abundance of all species *N_j_*.

Accordingly, for different outcomes for the same initial abundances of *N* species another variable has to be present, which in our case is the environment. The value of this environment could have been changed in the past by species activity, and thus, memory can occur.

#### 2) Separation of timescales between the change in the environment and population densities

The change of the environmental variable has to be rather slow compared to the change in species abundance. This can be seen from the main text Fig. 3c but also directly obtained from the underlying model:

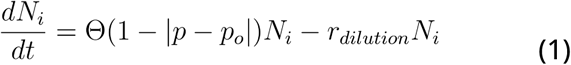

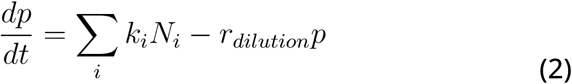

If the change of *p* is much faster than the change of *N_i_*, *p* can be considered in a steady state (dp/dt=0), and thus from Equation 2 we obtain

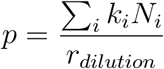

which gives with equation (1)

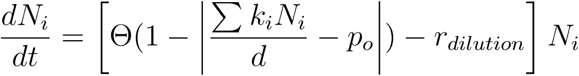

Accordingly, the change of *N_i_* does not depend on *p* anymore. Potential past changes of *p* will not affect the dynamics of *N_i_* and, thus, the interaction outcomes. That means memory has no impact anymore.

**Supplementary Fig 5:**
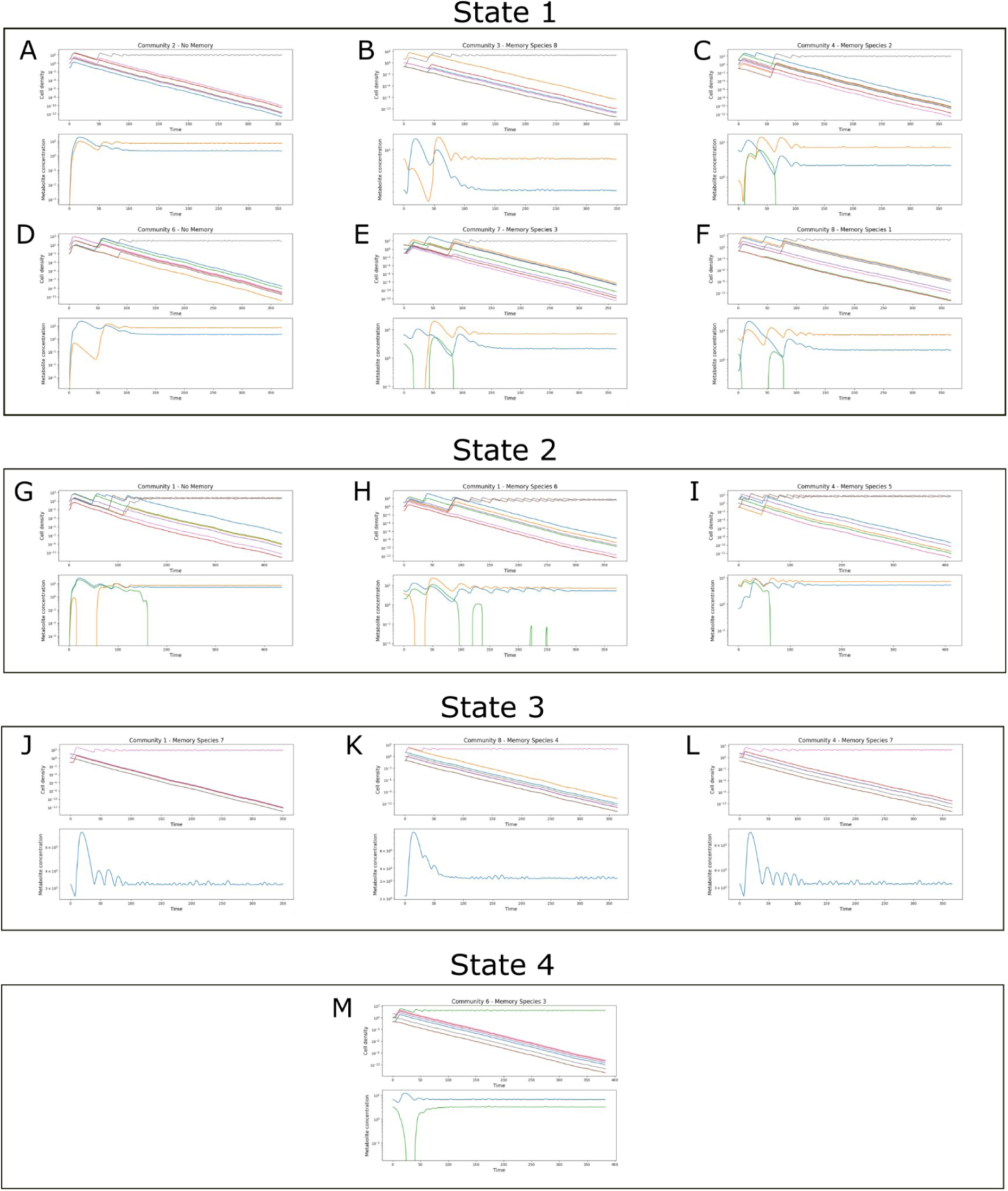
Example plots from simulations over equation 1 and 2, depicting cell density and metabolite concentration change over time. Bray-curtis dissimilarities were calculated from the final community composition, which in turn were used for non-metric multidimensional scaling as well as hierarchical clustering. Clustering revealed 4 clusters, which are hereby referred to as states. Different communities differ in their initial abundances and in absence of any memory may end up in different states, as observed in **A, G**. Identical communities can end up in different states in presence of different memories initially, as seen in **C, I, L** and in **H, J**.

**Supplementary Fig 6:**
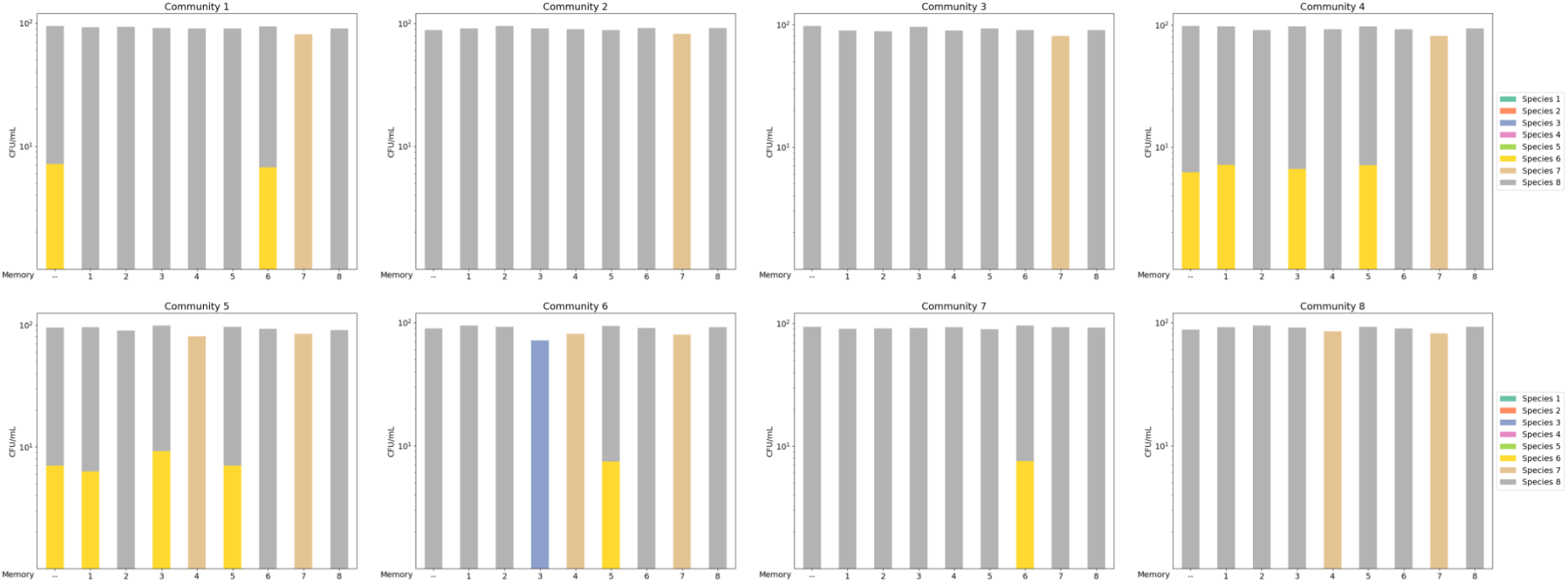
End-point relative abundances of 8 different communities, derived from simulations based on equation 1 and 2. Despite variations in their initial relative abundances, the communities are composed of the same eight species. Each plot represents sample communities that began with identical relative abundances but varied in their memories.

**Supplementary Fig 7:**
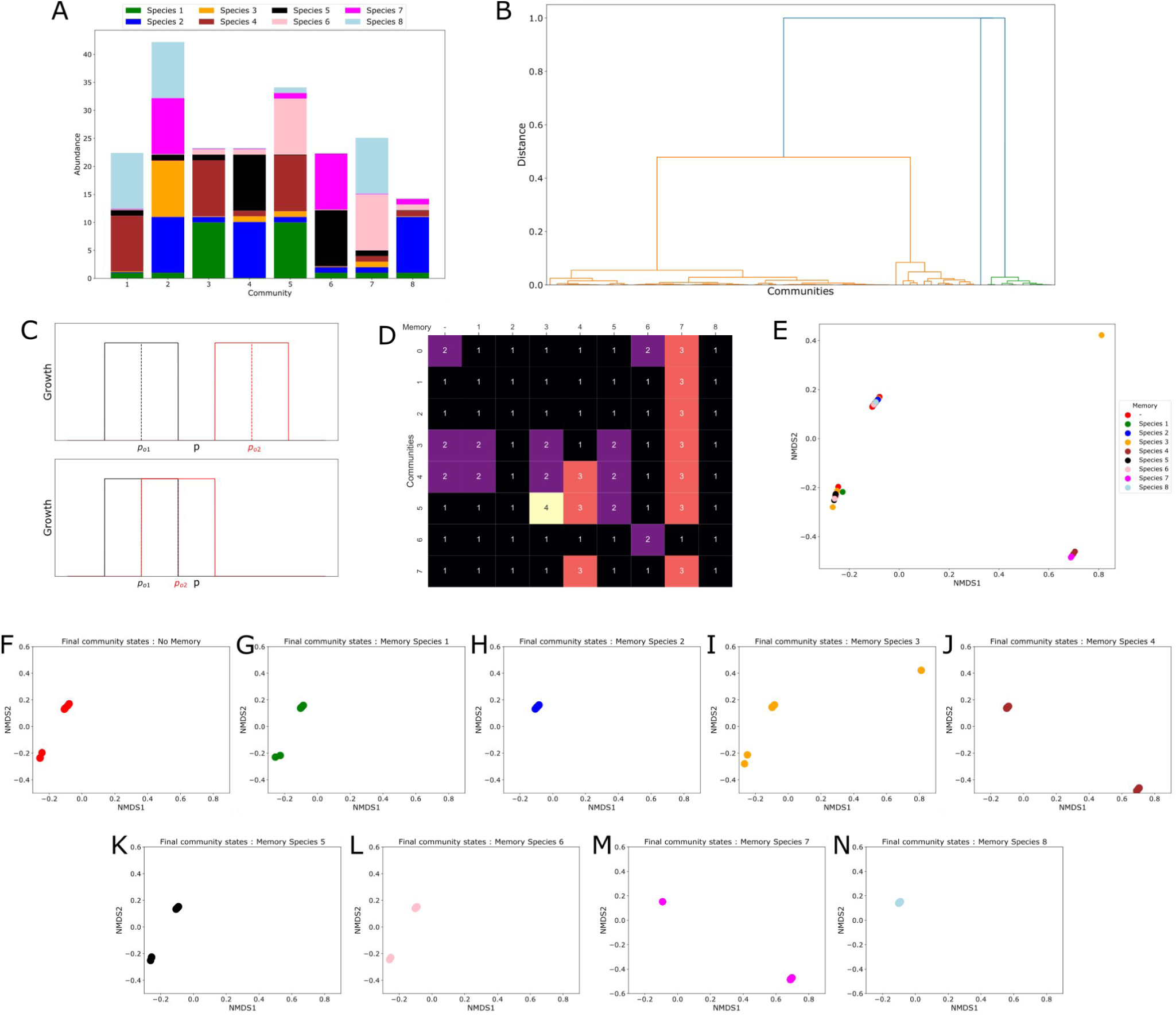
Simulation of first set of species using equations 1 and 2. **(A)** Initial composition of communities using first set of species. Each community is composed of 8 species, where each abundance is randomly selected from set [0.1,1,10]. **(B)** Hierarchical clustering on simulation derived-final community composition data across all initial compositions and supernatants reveal 4 clusters (also referred to as states) (**C)** Distribution of clusters/states in different memories across all communities reveals abundance of less frequent states in memories from certain species. State 2 was only present in communities assembled in presence of species 3 supernatant. (**C)** In order for ensure growth of both microbes, **optimum *p*** values of species within the community must be close to each other, such that their ‘growth zones’ coincide. (D) Heatmaps depicting States reached in presence of different memories across all communities. State 3 is primarily achieved in presence of Species 7 memory, whilst communities can end up in different states in absence of memory. (**E)** Similarly, nMDS plot based on Bray-curtis dissimilarities between final community compositions across all communities shows clustering into 4 distinct clusters, which are similar to those in Hierarchical clustering. **(F), (G), (H), (I), (J), (K), (L), (M), (N)** Single memory (or absence thereof) nMDS plots from **(E).**

**Supplementary Fig 8:**
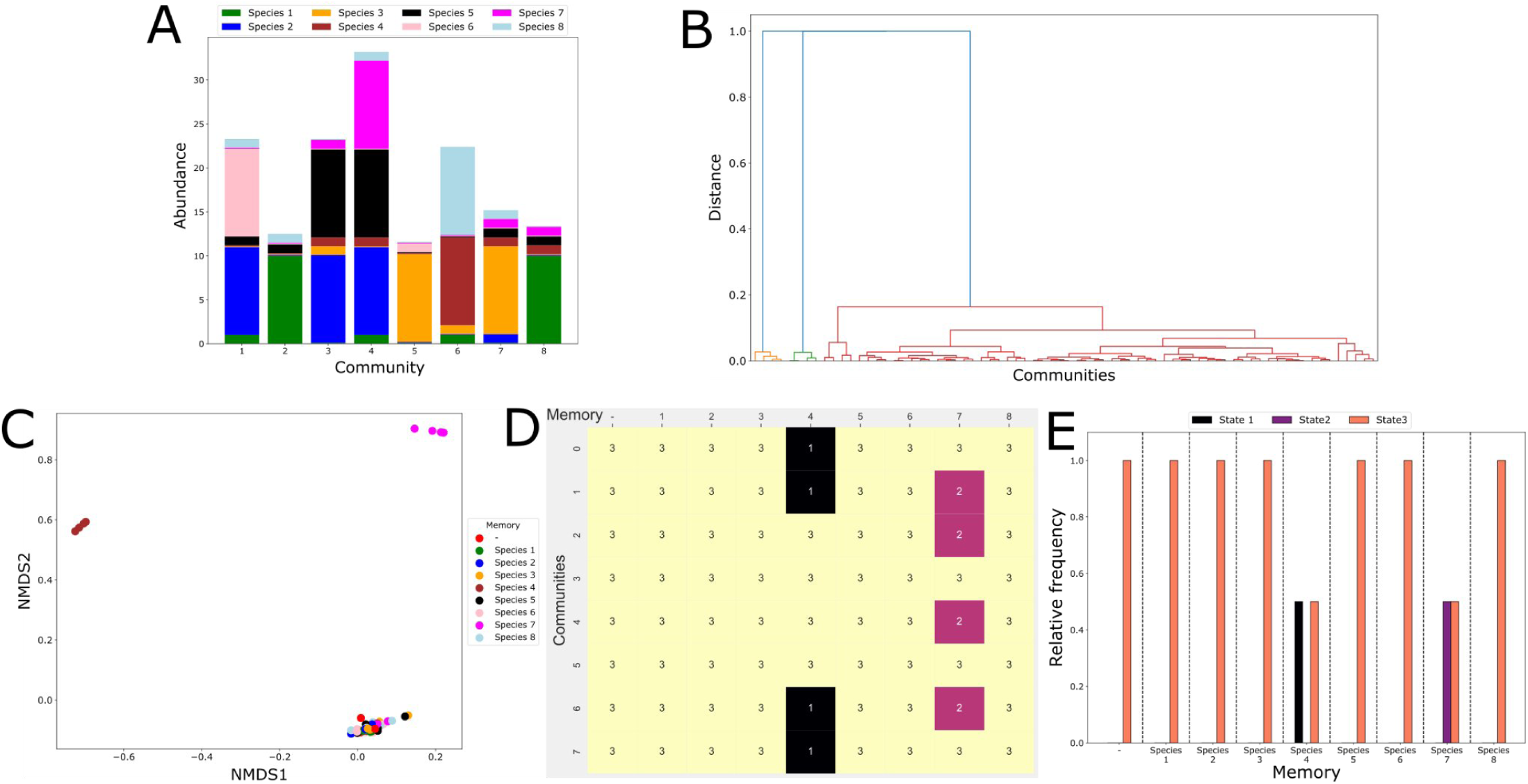
Second set of simulations. **(A)** Initial composition of communities using second set of species. Each community is composed of 8 species, where each abundance is randomly selected from [0.1,1,10] set. **(B)** Post simulation, Bray-curtis dissimilarities were computed on end-point community composition and Hierarchical clustering reveals 3 major clusters (or states). **(C)** nMDS projection of Bray-Curtis dissimilarities between end-state communities also points out at 3 cluster. Cluster ids are similar as compared to hierarchical clustering. **(D)** Heatmap of states achieved in different communities in presence of memories from individual constituent species. **(E)** Presence of memories from either species 4 or species 7 leads to increase in frequency of community ending up in either state 1 or 2.

**Supplementary Fig. 9:**
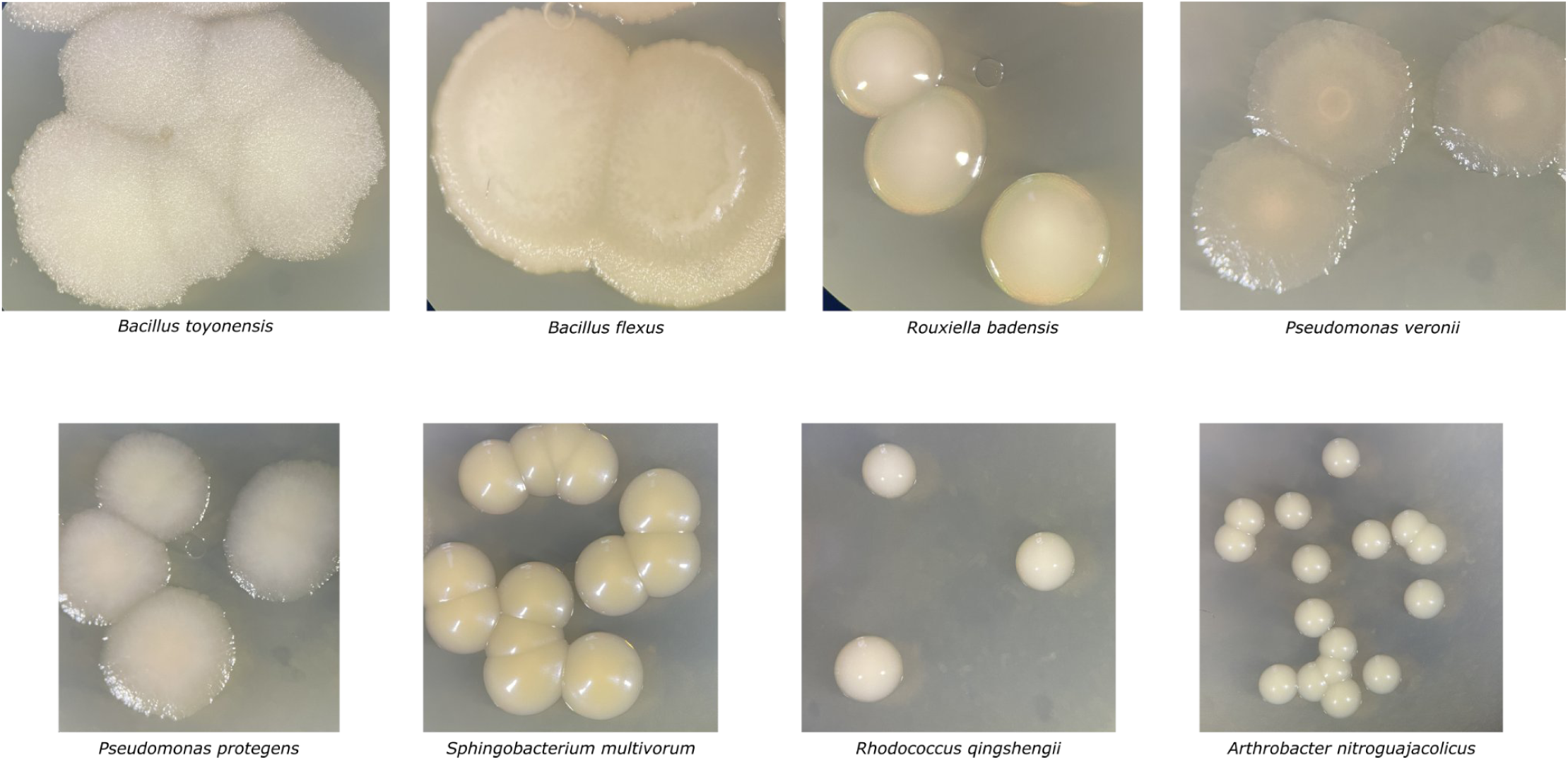
Colony morphologies of 8 *C. elegans* gut species. Each species was identified using size, color and specific distinct features. *Bacillus toyonensis* displays ‘ white, granular and rough’ surface, while *Bacillus flexus* was identified as ‘large, yellow and rough’ colonies. *R. qingshengii* appears as ‘small, white and shiny’ colonies while *A. nitroguajacolicus* appears as ‘small, greenish and shiny colonies’. Similarly other species were identifiable with distinct features.

**Supplementary Fig 10:**
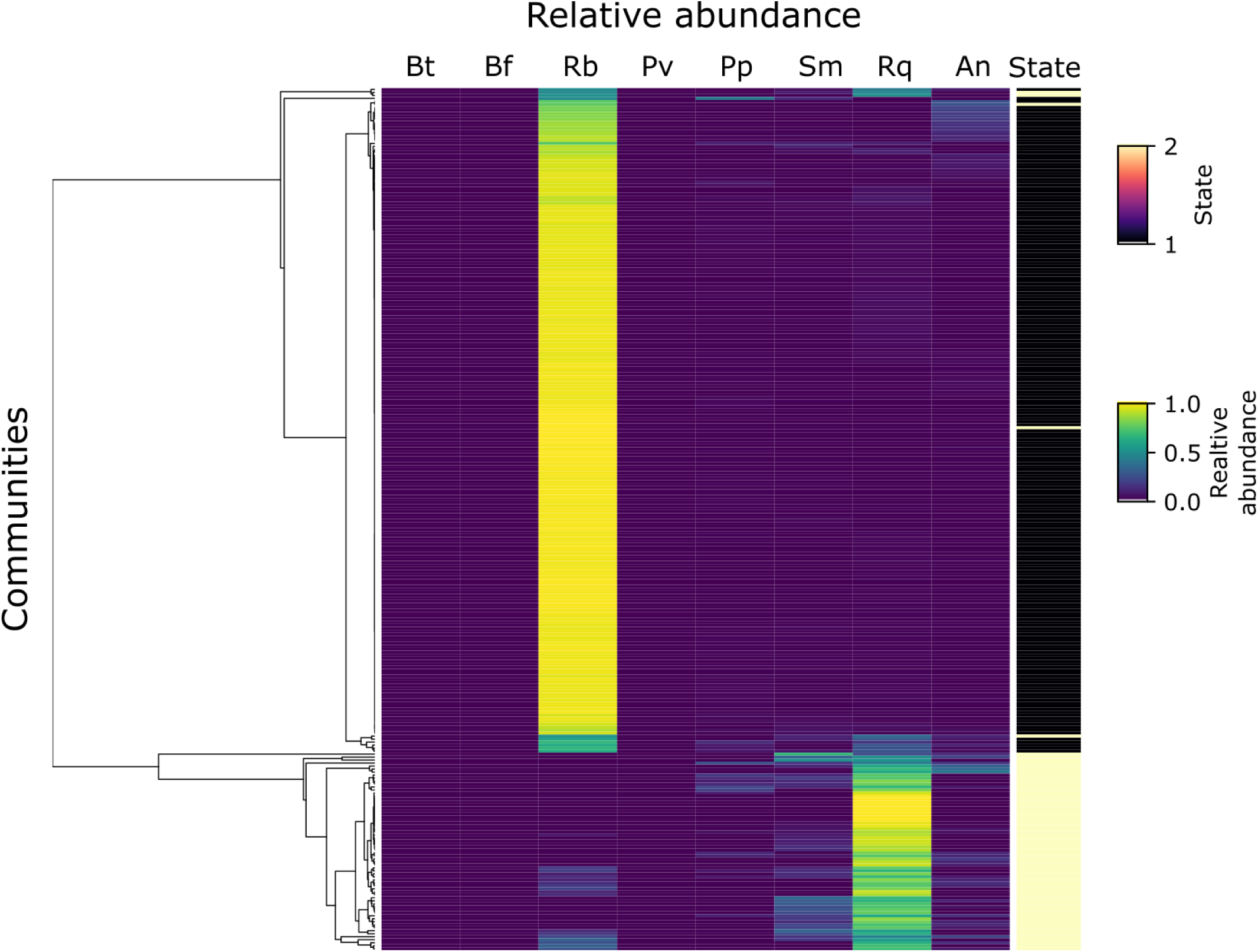
Complex heatmap representing relative abundances of each species in the final community compositions (each row represents one community replicate). Clustering based on relative abundances reveals State 1 communities is mostly dominated by ***Rb***, while State 2 communities exhibit presence of ***Rq, Sm, An, Pp*** and ***Rb*.**

**Supplementary Fig 11:**
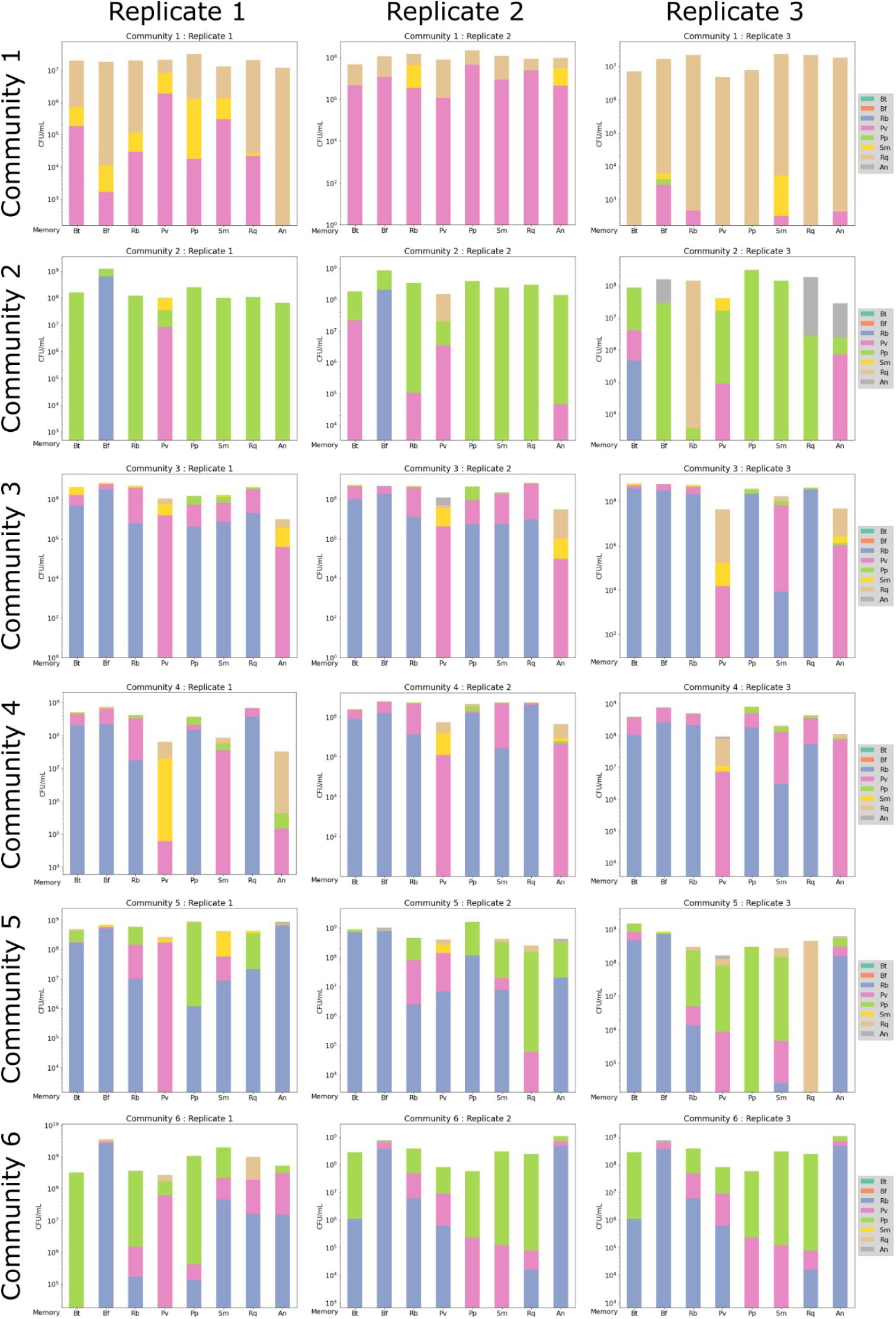
Community assembly experiment on Day 5. Community composition on Day 5, determined from plating on NM agar. Each row shows a different initial community composition. Columns corresponds technical replicates. The bars in each plot show the development of initial communities in the population memory of the species named on the x-axis. Within each bar, species are stacked linearly, while bars in bar plots are log-scaled, thereby maintaining relative distribution of species.

**Supplementary Fig 12:**
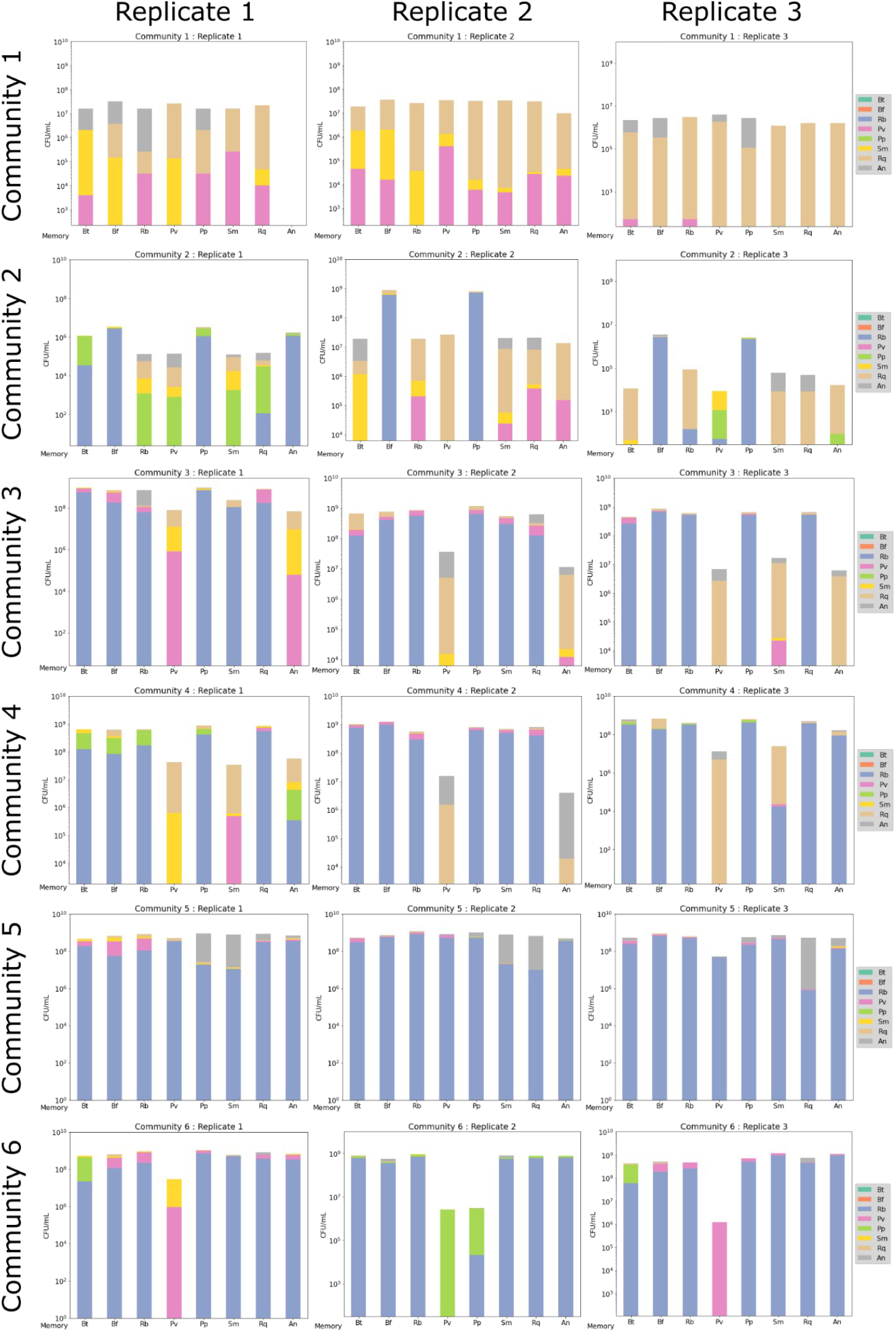
Community assembly experiment on Day 10. Community composition on Day 10, determined from plating on NM agar. Each row shows a different initial community composition. Columns corresponds technical replicates. The bars in each plot show the development of initial communities in the population memory of the species named on the x-axis. Within each bar, species are stacked linearly, while bars in bar plots are log-scaled, thereby maintaining relative distribution of species.

**Supplementary Fig 13:**
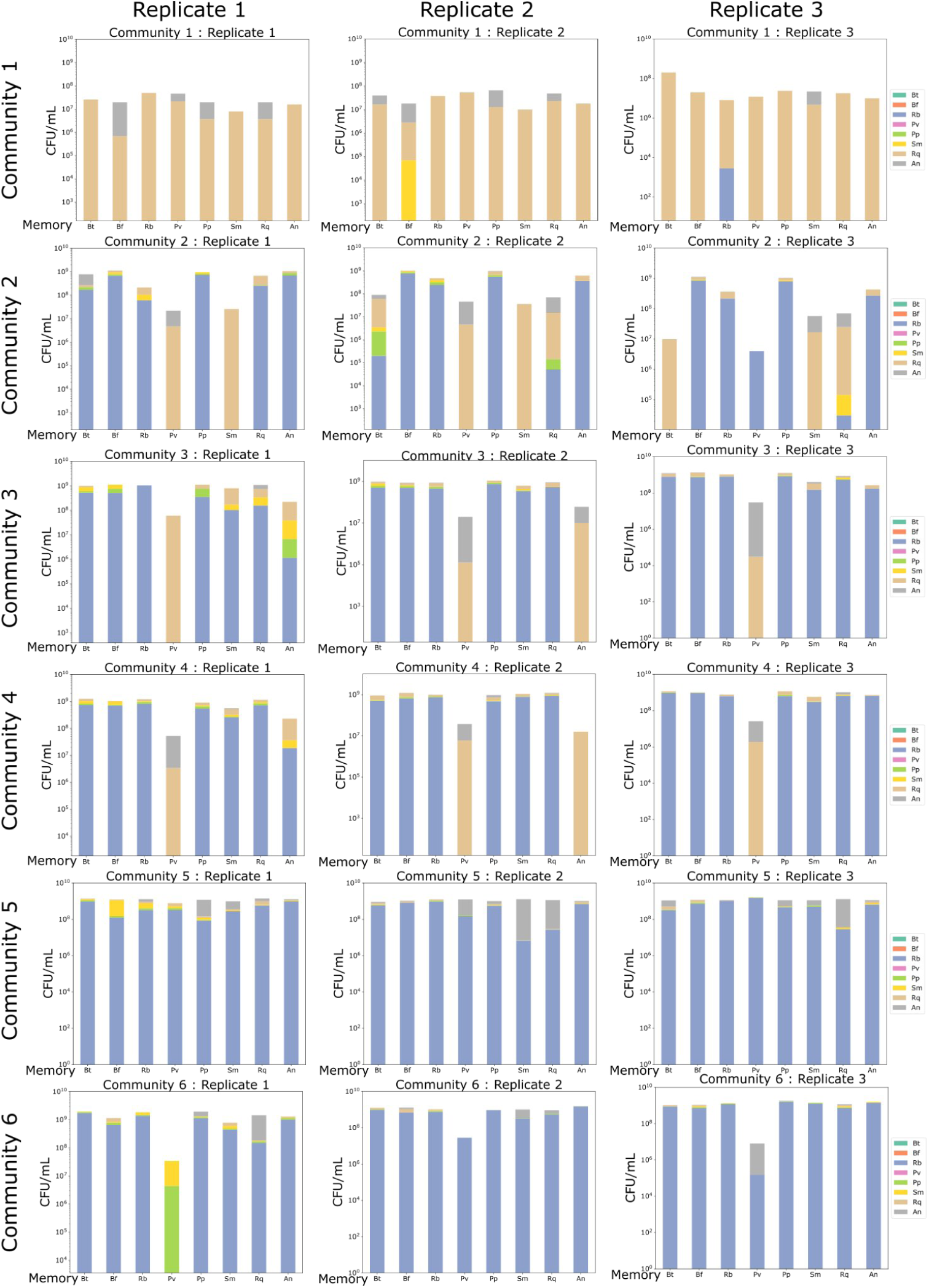
Community assembly experiment on Day 12. Community composition on Day 12, determined from plating on NM agar. Each row shows a different initial community composition. Columns corresponds technical replicates. The bars in each plot show the development of initial communities in the population memory of the species named on the x-axis. Within each bar, species are stacked linearly, while bars in bar plots are log-scaled, thereby maintaining relative distribution of species.

**Supplementary Fig 14:**
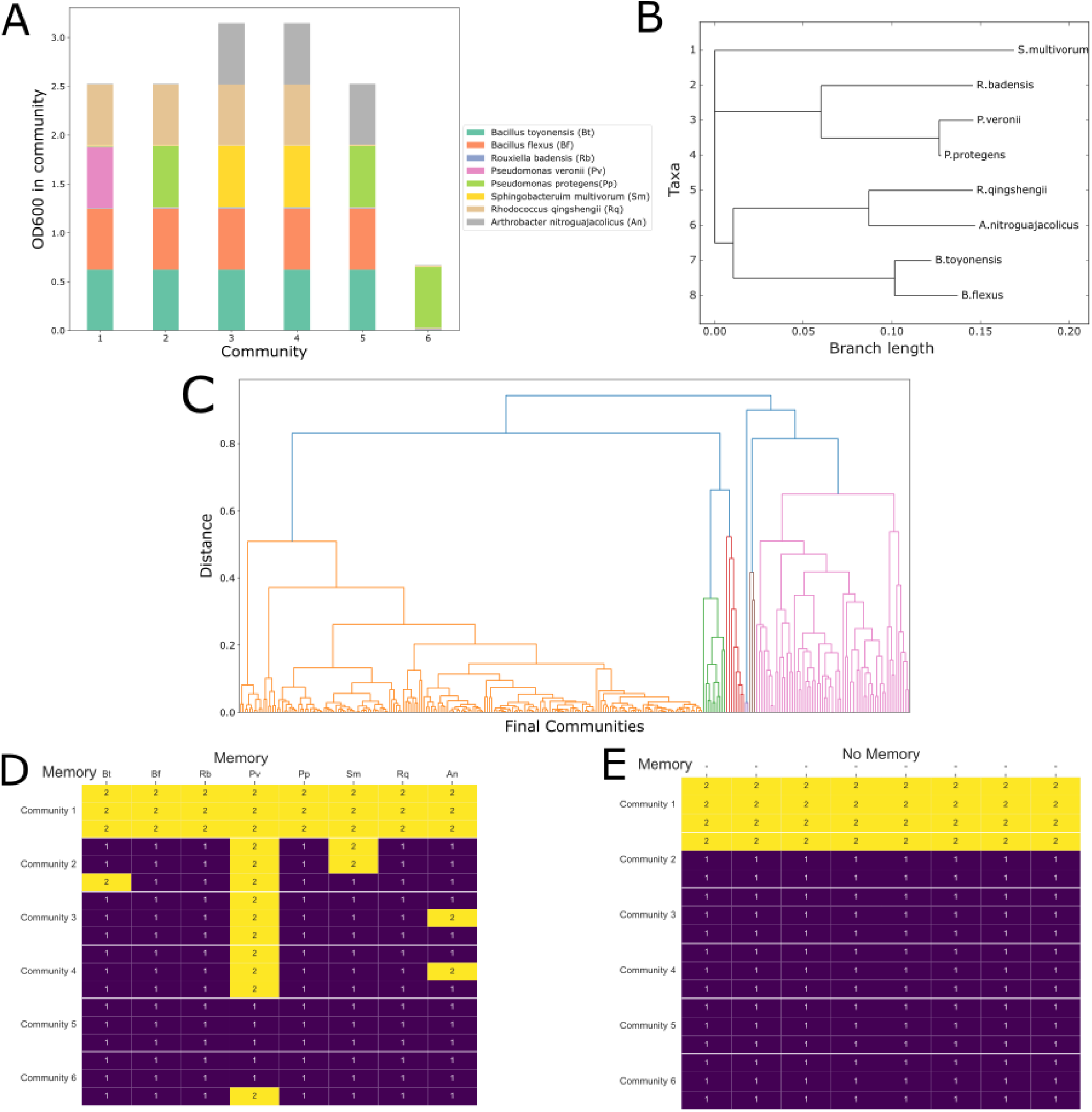
**A.** Final OD600 values of microbial species in initial community assembly. Equivolume proportions of each species were assembled either at high (OD600 = 5) or low (OD600 = 0.05) ODs into 6 different communities. Each community was assembled with same 8 species, but with different initial proportions. **B.** Bacterial species used in community assembly represent diverse taxa as noted in Phylogenetic tree. **C.** Final community composition was obtained by plating at Day 12 and Bray-curtis dissimilarities were computed. Hierarchical clustering based on dissimilarities revealed 2 major clusters, and cluster identities were used to generate heatmaps **D, E.** X-axis represents memories of different species and on Y-axis are different communities. Horizontal white lines separate 3 technical replicates of each community. **D.** Memory of *Pseudomonas veronii* leads to communities ending up in State 2 more frequently as compared to memories of other species, **E.** as well as communities assembled in absence of supernatant.

**Supplementary Figure 15:**
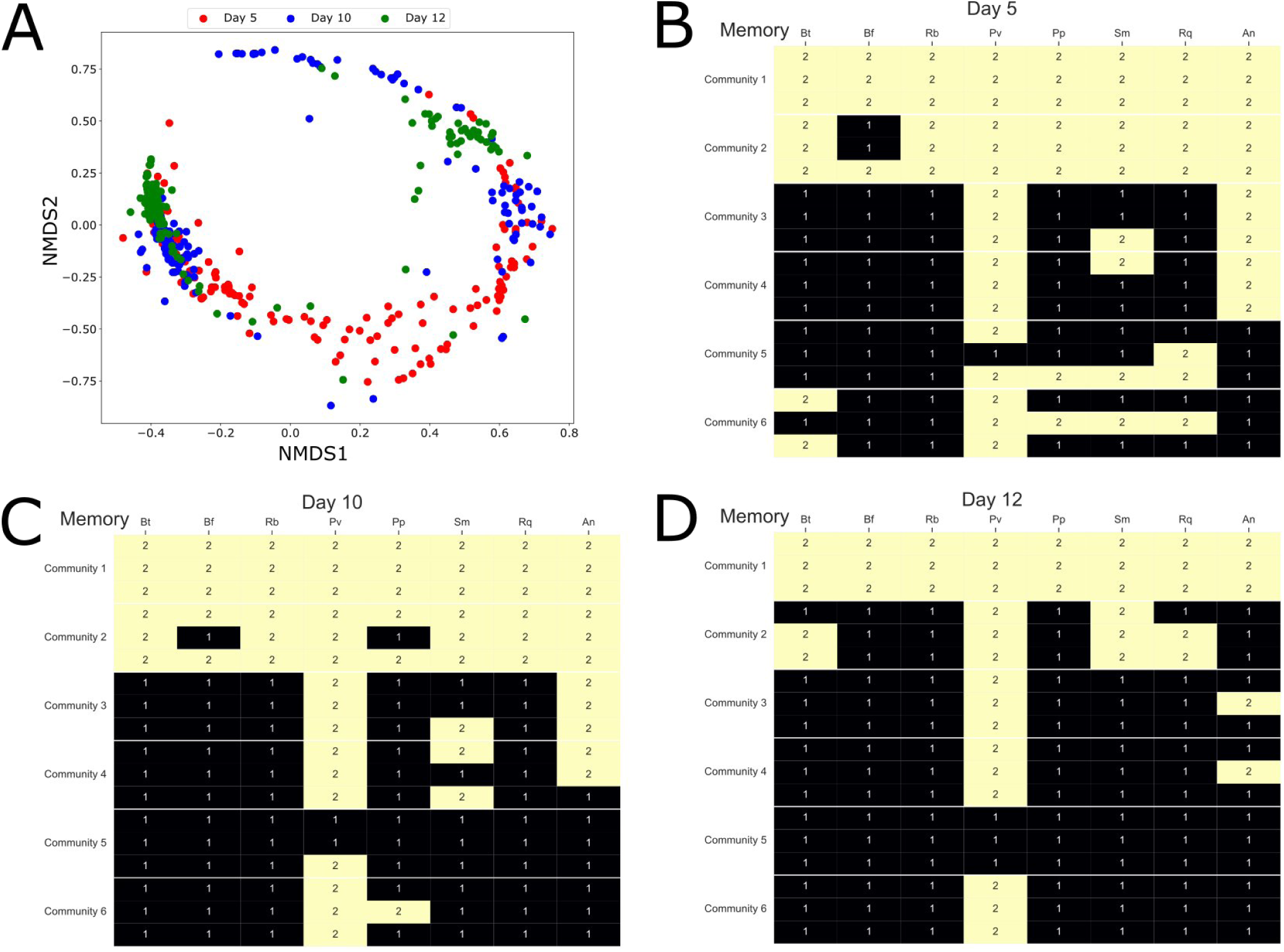
**(A)** To study distribution and transitioning of states, community composition data of only communities assembled in presence of memories from Day 5, Day 10 and Day 12 was combined and NMDS plot were computed from Bray-curtis dissimilarities. Distribution of samples within NMDS plot reveals more dispersed distribution of Day 5 samples, which appear to be converging on Day 10 towards tightly clustered State 1 and 2 on Day 12. **(B), (C), (D)** Communities assembled in memory of *Pseudomonas veronii (Pv),* which end up in State 2, mostly continue to remain in State 2 as observed in State heatmaps.

## References

1. Kandel, E. R., Dudai, Y. & Mayford, M. R. The Molecular and Systems Biology of Memory. Cell 157, 163–186 (2014).

2. Henikoff, S. & Greally, J. M. Epigenetics, cellular memory and gene regulation. Curr. Biol. 26, R644–R648 (2016).

3. Turner, B. M. Cellular Memory and the Histone Code. Cell 111, 285–291 (2002).

4. Acar, M., Becskei, A. & van Oudenaarden, A. Enhancement of cellular memory by reducing stochastic transitions. Nature 435, 228–232 (2005).

5. Guan, Q., Haroon, S., Bravo, D. G., Will, J. L. & Gasch, A. P. Cellular Memory of Acquired Stress Resistance in Saccharomyces cerevisiae. Genetics 192, 495–505 (2012).

6. Norman, T. M., Lord, N. D., Paulsson, J. & Losick, R. Memory and modularity in cell-fate decision making. Nature 503, 481–486 (2013).

7. Mutlu, A. et al. Phenotypic memory in Bacillus subtilis links dormancy entry and exit by a spore quantity-quality tradeoff. Nat. Commun. 9, 69 (2018).

8. Miyaue, S. et al. Bacterial Memory of Persisters: Bacterial Persister Cells Can Retain Their Phenotype for Days or Weeks After Withdrawal From Colony–Biofilm Culture. Front. Microbiol. 9, (2018).

9. Jackson, D. E., Martin, S. J., Holcombe, M. & Ratnieks, F. L. W. Longevity and detection of persistent foraging trails in Pharaoh’s ants, Monomorium pharaonis (L.). Anim. Behav. 71, 351–359 (2006).

10. Reid, C. R., Latty, T., Dussutour, A. & Beekmani, M. Slime mold uses an externalized spatial “memory” to navigate in complex environments. Proc. Natl. Acad. Sci. 109, 17490–17494 (2012).

11. Smith-Ferguson, J. & Beekman, M. Who needs a brain? Slime moulds, behavioural ecology and minimal cognition. Adapt. Behav. 28, 465–478 (2020).

12. Douglas, A. E. The microbial exometabolome: ecological resource and architect of microbial communities. Philos. Trans. R. Soc. B Biol. Sci. 375, 20190250 (2020).

13. Estrela, S. et al. Environmentally Mediated Social Dilemmas. Trends Ecol. Evol. 34, 6–18 (2019).

14. Ratzke, C. & Gore, J. Modifying and reacting to the environmental pH can drive bacterial interactions. PLOS Biol. 16, e2004248 (2018).

15. Østlie, H. M., Treimo, J. & Narvhus, J. A. Effect of temperature on growth and metabolism of probiotic bacteria in milk. Int. Dairy J. 15, 989–997 (2005).

16. Vollbrecht, D. & Schlegel, H. G. Excretion of metabolites by hydrogen bacteria II. Influences of aeration, pH, temperature, and age of cells. Eur. J. Appl. Microbiol. Biotechnol. 6, 157–166 (1978).

17. Hoerr, V. et al. Characterization and prediction of the mechanism of action of antibiotics through NMR metabolomics. BMC Microbiol. 16, 82 (2016).

18. Smith, R. et al. Programmed Allee effect in bacteria causes a tradeoff between population spread and survival. Proc. Natl. Acad. Sci. U. S. A. 111, 1969–1974 (2014).

19. Ratzke, C. & Gore, J. Self-organized patchiness facilitates survival in a cooperatively growing Bacillus subtilis population. Nat. Microbiol. 1, 1–5 (2016).

20. Ratzke, C. & Gore, J. Modifying and reacting to the environmental pH can drive bacterial interactions. PLOS Biol. 16, e2004248 (2018).

21. Clarke, A. & Fraser, K. P. P. Why does metabolism scale with temperature? Funct. Ecol. 18, 243–251 (2004).

22. Smith, T. P. et al. Community-level respiration of prokaryotic microbes may rise with global warming. Nat. Commun. 10, 5124 (2019).

23. Østlie, H. M., Treimo, J. & Narvhus, J. A. Effect of temperature on growth and metabolism of probiotic bacteria in milk. Int. Dairy J. 15, 989–997 (2005).

24. Murphy, M. G. & Condon, S. Comparison of aerobic and anaerobic growth of Lactobacillus plantarum in a glucose medium. Arch. Microbiol. 138, 49–53 (1984).

25. Clarke, K. R. Non-parametric multivariate analyses of changes in community structure. Aust. J. Ecol. 18, 117–143 (1993).

26. Zhang, C., Kong, Y., Xiang, Q., Ma, Y. & Guo, Q. Bacterial memory in antibiotic resistance evolution and nanotechnology in evolutionary biology. iScience 26, 107433 (2023).

27. Casadesús, J. & D’Ari, R. Memory in bacteria and phage. BioEssays 24, 512–518 (2002).

28. Bhattacharyya, S., et al. A heritable iron memory enables decision-making in Escherichia coli. Proc. Natl. Acad. Sci. 120, e2309082120 (2023).

29. Ronin, I., Katsowich, N., Rosenshine, I. & Balaban, N. Q. A long-term epigenetic memory switch controls bacterial virulence bimodality. eLife 6, e19599 (2017).

30. Wolf, D. M. et al. Memory in Microbes: Quantifying History-Dependent Behavior in a Bacterium. PLOS ONE 3, e1700 (2008).

31. Lambert, G. & Kussell, E. Memory and Fitness Optimization of Bacteria under Fluctuating Environments. PLOS Genet. 10, e1004556 (2014).

32. Gokhale, C. S., Giaimo, S. & Remigi, P. Memory shapes microbial populations. PLOS Comput. Biol. 17, e1009431 (2021).

33. Mathis, R. & Ackermann, M. Asymmetric cellular memory in bacteria exposed to antibiotics. BMC Evol. Biol. 17, 73 (2017).

34. Kordes, A. et al. Establishment of an induced memory response in Pseudomonas aeruginosa during infection of a eukaryotic host. ISME J. 13, 2018–2030 (2019).

35. Gardner, T. S., Cantor, C. R. & Collins, J. J. Construction of a genetic toggle switch in Escherichia coli. Nature 403, 339–342 (2000).

36. Vermeersch, L. et al. Do microbes have a memory? History-dependent behavior in the adaptation to variable environments. Front. Microbiol. 13, (2022).

37. Miyaue, S. et al. Bacterial Memory of Persisters: Bacterial Persister Cells Can Retain Their Phenotype for Days or Weeks After Withdrawal From Colony–Biofilm Culture. Front. Microbiol. 9, (2018).

38. Martino, R. D., Picot, A. & Mitri, S. Oxidative stress changes interactions between 2 bacterial species from competitive to facilitative. PLOS Biol. 22, e3002482 (2024).

39. Ratzke, C. & Gore, J. Modifying and reacting to the environmental pH can drive bacterial interactions. PLOS Biol. 16, e2004248 (2018).

40. Goldford, J. E. et al. Emergent simplicity in microbial community assembly. Science 361, 469–474 (2018).

41. West, S. A., Diggle, S. P., Buckling, A., Gardner, A. & Griffin, A. S. The Social Lives of Microbes. Annu. Rev. Ecol. Evol. Syst. 38, 53–77 (2007).

42. Harrison, F., Paul, J., Massey, R. C. & Buckling, A. Interspecific competition and siderophore-mediated cooperation in Pseudomonas aeruginosa. ISME J. 2, 49–55 (2008).

43. Miller, M. B. & Bassler, B. L. Quorum Sensing in Bacteria. Annu. Rev. Microbiol. 55, 165– 199 (2001).

44. Stubbendieck, R. M., Vargas-Bautista, C. & Straight, P. D. Bacterial Communities: Interactions to Scale. Front. Microbiol. 7, (2016).

45. Couzin, I. D., Krause, J., James, R., Ruxton, G. D. & Franks, N. R. Collective Memory and Spatial Sorting in Animal Groups. J. Theor. Biol. 218, 1–11 (2002).

46. Jackson, D. E. & Ratnieks, F. L. W. Communication in ants. Curr. Biol. 16, R570–R574 (2006).

47. Debray, R. et al. Priority effects in microbiome assembly. Nat. Rev. Microbiol. 20, 109–121 (2022).

48. Carlström, C. I. et al. Synthetic microbiota reveal priority effects and keystone strains in the Arabidopsis phyllosphere. *Nat*. Ecol. Evol. 3, 1445–1454 (2019).

49. A strong priority effect in the assembly of a specialized insect-microbe symbiosis.

50. Ogle, K. et al. Quantifying ecological memory in plant and ecosystem processes. Ecol. Lett. 18, 221–235 (2015).

51. Schweiger, A. H., Boulangeat, I., Conradi, T., Davis, M. & Svenning, J.-C. The importance of ecological memory for trophic rewilding as an ecosystem restoration approach. Biol. Rev. 94, 1–15 (2019).

52. Webster, C. R. et al. Promoting and maintaining diversity in contemporary hardwood forests: Confronting contemporary drivers of change and the loss of ecological memory. For. Ecol. Manag. 421, 98–108 (2018).

53. Odling-Smee, F. J., Laland, K. N. & Feldman, M. W. Niche Construction. Am. Nat. 147, 641–648 (1996).

54. Day, R. L., Laland, K. N. & Odling-Smee, F. J. Rethinking Adaptation: The Niche-Construction Perspective. Perspect. Biol. Med. 46, 80–95 (2003).

55. Estrela, S., Diaz-Colunga, J., Vila, J. C. C. & Sanchez-Gorostiaga, A. Diversity begets diversity under microbial niche construction.

56. https://www.biorxiv.org/content/10.1101/2023.10.31.565024v1.full.pdf. https://www.biorxiv.org/content/10.1101/2023.10.31.565024v1.full.pdf.

57. Quintus, S. & Allen, M. S. Niche Construction and Long-Term Trajectories of Food Production. J. Archaeol. Res. 32, 209–261 (2024).

58. Boivin, N. L., et al. Ecological consequences of human niche construction: Examining long-term anthropogenic shaping of global species distributions. Proc. Natl. Acad. Sci. 113, 6388–6396 (2016).

59. van den Berg, N. I., et al. Ecological modelling approaches for predicting emergent properties in microbial communities. *Nat*. Ecol. Evol. 6, 855–865 (2022).

60. Friedman, J., Higgins, L. M. & Gore, J. Community structure follows simple assembly rules in microbial microcosms. *Nat*. Ecol. Evol. 1, 1–7 (2017).

61. Stein, R. R. et al. Ecological Modeling from Time-Series Inference: Insight into Dynamics and Stability of Intestinal Microbiota. PLOS Comput. Biol. 9, e1003388 (2013).

62. Kuntal, B. K., Gadgil, C. & Mande, S. S. Web-gLV: A Web Based Platform for Lotka-Volterra Based Modeling and Simulation of Microbial Populations. Front. Microbiol. 10, (2019).

63. Momeni, B., Xie, L. & Shou, W. Lotka-Volterra pairwise modeling fails to capture diverse pairwise microbial interactions. eLife 6, e25051 (2017).

64. Ratzke, C., Denk, J. & Gore, J. Ecological suicide in microbes. *Nat*. Ecol. Evol. 2, 867–872 (2018).

65. Gomaa, E. Z. Human gut microbiota/microbiome in health and diseases: a review. Antonie Van Leeuwenhoek 113, 2019–2040 (2020).

66. Manor, O. et al. Health and disease markers correlate with gut microbiome composition across thousands of people. Nat. Commun. 11, 5206 (2020).

67. Hagan, T. et al. Antibiotics-Driven Gut Microbiome Perturbation Alters Immunity to Vaccines in Humans. Cell 178, 1313–1328.e13 (2019).

68. Letourneau, J. et al. Ecological memory of prior nutrient exposure in the human gut microbiome. ISME J. 1–12 (2022) doi:10.1038/s41396-022-01292-x.

69. Khazaei, T. et al. Metabolic multistability and hysteresis in a model aerobe-anaerobe microbiome community. Sci. Adv. 6, eaba0353 (2020).

70. Zorgani, A. & Das, B. C. Exploring the memory of the gut microbiome: a multifaceted perspective. Front. Microbiomes 3, (2024).

71. Stacy, A. et al. Infection trains the host for microbiota-enhanced resistance to pathogens. Cell 184, 615–627.e17 (2021).

72. Ratajczak, W. et al. Immunomodulatory potential of gut microbiome-derived short-chain fatty acids (SCFAs). Acta Biochim. Pol. (2019) doi:10.18388/abp.2018_2648.

73. Henrick, B. M. et al. Elevated Fecal pH Indicates a Profound Change in the Breastfed Infant Gut Microbiome Due to Reduction of *Bifidobacterium* over the Past Century. mSphere 3, e00041–18 (2018).

